# Scalable, fast and accurate differential gene expression testing from millions of cells of multiple patients

**DOI:** 10.1101/2025.07.24.666556

**Authors:** Giovanni Santacatterina, Niccolò Tosato, Salvatore Milite, Katsiaryna Davydzenka, Edoardo Insaghi, Guido Sanguinetti, Stefano Cozzini, Leonardo Egidi, Giulio Caravagna

**Author notes:** These authors contributed equally (Giovanni Santacatterina, Niccolò Tosato). These authors jointly supervised this work (Stefano Cozzini, Leonardo Egidi, Giulio Caravagna).

## Abstract

Since the development of DNA microarrays and later RNA bulk sequencing, testing with statistically independent samples has been the standard method for detecting genes with different transcription patterns. Single-cell assays challenge these assumptions because individual cells are statistically dependent, and all proposed methodologies present mathematical limitations or computational bottlenecks that prevent a seamless integration of data from many cells and patients simultaneously. In this work, we solve this crucial limitation by introducing a Bayesian framework that retrieves the independence structure at the level of individual patients, separating differences across individuals from actual transcriptional differences. Leveraging multi-GPU and variational inference, our approach excels across different experimental designs and scales to analyse over 10 million cells. This framework enables single-cell differential expression analysis that can finally integrate datasets from large clinical cohorts, atlas projects, or drug-response screens with thousands of samples and millions of cells.

## Introduction

Since the advent of DNA microarrays in the late 1990s^1^, statistical testing for differential gene expression has played a central role in the discovery of condition-specific transcriptional programs^2^. Classical approaches, based on linear models and statistical tests for the estimated parameters, assume that expression measurements across biological replicates are statistically independent. These assumptions held in early bulk RNA sequencing studies, where each sample corresponded to a distinct individual or experimental unit, and the field of differential expression (DE) testing has flourished over the years. Today, specific tools such as DeSeq2^3^, edgeR^4^, and limma^5^ are virtually used in every article that performs bulk RNA sequencing.

However, with the growing use of single-cell RNA sequencing (scRNA-seq), these fundamental assumptions have been called into question^6^. In single-cell assays, we obtain tens of thousands of gene expression measurements from individual cells within a single biological replicate. However, these cells are not independent observations; they share a lineage, exposure to the same micro-environment, and often technical artefacts that depend on the sequencing technology^7,8^. Treating cells as independent replicates can overstate statistical significance (type-I errors) in single-cell differential expression (scDE) studies, an issue especially pronounced in multi-sample designs where the main source of biological replication is at the subject level^6^. Therefore, the field has established, see “Challenge X: Integration of single-cell data across samples, experiments, and types of measurement” by Lähnemann et al. ^6^, that we need to develop statistically robust scDE tools that account for this hierarchical structure^9,10^, distinguishing within-patient (i.e., cell-to-cell heterogeneity) from between-patient variability (Fig.1a,b). This is crucial because the latter confounds population-level inferences when we compute p-values and log-fold change (LFC) statistics.

Many different strategies have been adopted to address this challenge (Supplementary Table S1), drawing from batch-effect removal and model-based scDE, each with distinct advantages and limitations^3–5,11–14^. Batch-integration approaches (e.g., ComBat^11^ and Harmony^12^) can be used to treat patient identity as a technical batch effect to be harmonized. These methods may struggle to preserve biologically meaningful inter-patient variation and can underperform compared to simpler covariate modeling^15^. Instead, model-based scDE approaches explicitly partition variance sources, building pseudobulk, generalized or mixed models that simultaneously account for within– and between-patient variability. Pseudobulk aggregation strategies combine counts per patient, then apply tools established for bulk RNA sequencing (e.g., limma or DESeq2). While this approach is the fastest, it obscures cell-to-cell variability, potentially missing subtle but biologically relevant differential signals. This issue is commonly addressed by fitting generalized linear models (GLMs) with patient-specific fixed effects in the design matrix (e.g., using glmGamPoi^13^). While such models are statistically valid, they can become cumbersome in practice as the number of patients increases, especially when patient identity is closely aligned with experimental conditions. In these settings, fixed-effect parametrizations increase model complexity and computational cost, motivating alternative approaches that account for patient-level variability without explicitly estimating many patient-specific effects. Finally, mixed models (e.g., NEBULA^14^) fit a GLM with patients as a random effect. This approach demonstrates better control of false positives and higher power than baseline linear models. Still, it introduces patient-specific parameters that limit the scalability of this approach to a limited number of individuals. Therefore, significant challenges remain in scaling these models to population-level studies that can include data from thousands of patients and measurements, virtually scaling beyond several million data points.

Here we present a scalable and statistically sound framework, called DEVIL, designed to address said challenges in the context of scDE testing. Our model involves a Bayesian Gamma-Poisson structure, like glmGamPoi^13^, to capture variability at both the gene and cell levels. Additionally, our model exploits clustered sandwich estimators^16,17^, with clusters defined by patient identity, to produce standard errors that are robust to within-patient correlation structure, without requiring that structure to be explicitly modeled. In practical terms, our approach is a GLM that captures correlations within each patient without increasing the number of parameters, unlike mixed models. We achieve this using the sandwich estimators by computing an effective sample size that reflects the true degree of independent information in the data, which is substantially lower than the raw cell count, and using this to calibrate inference appropriately. In contrast to competing methodologies, our approach demonstrates generalizability across multiple experimental designs while scaling to over 10^7^ cells with thousands of genes. This is also achieved by a parallel variational inference algorithm tailored for modern multi-GPU architectures, enabling scDE analyses that can integrate scRNA-seq datasets from large clinical cohorts, atlas projects, or drug-response screens.

## Results

### Robust and efficient scDE testing for large-scale multi-patient designs

Canonical scDE testing from scRNA-seq data requires integrating measurements across multiple patients (population-level analysis), where cells are clustered within individuals and naïve independence assumptions inflate type-I error rates. We introduce DEVIL (differential expression with variational inference learning), a framework that enables both single-patient and population-level scDE analysis across millions of cells while accounting for this hierarchical structure (Fig. 1c,d). Like pseudobulk and other GLM-based methods, DEVIL assumes independence between patients while explicitly modelling the dependence structure within them.

**Figure 1.**
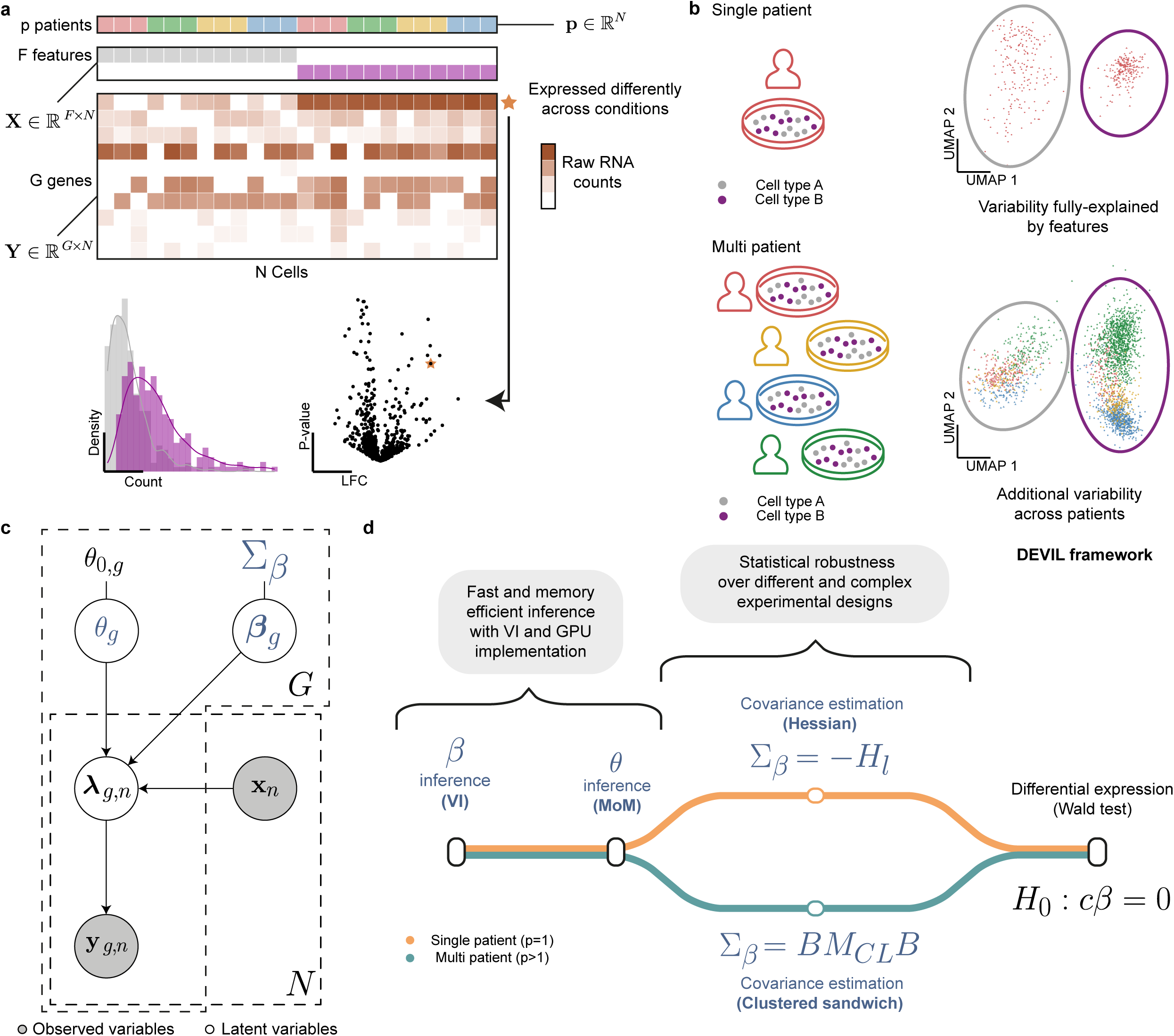
Multi-patient single-cell differential expression (scDE) testing with DEVIL. **a.** DEVIL operates on three primary inputs: **p** denoting patient assignments for each cell, design matrix **X**, encoding experimental covariates, and matrix **Y**, containing raw gene expression counts. Gene-level variation is illustrated through a histogram of expression values and a representative volcano plot, which shows log-fold change versus p-values, highlighting scDE patterns. **b.** Inter-patient expression heterogeneity challenges scDE testing. UMAP projections illustrate the contrast between single-patient and multi-patient datasets. While single-patient embeddings predominantly reflect intrinsic biological structure (e.g., cell type), multi-patient data introduces additional variation driven by patient-specific factors, underscoring the need for models that disentangle technical and biological sources of variation. **c.** The probabilistic model underlying DEVIL is based on a Gamma-Poisson framework that captures gene-specific overdispersion and population-level effects. Central parameters include gene-wise regression coefficients (***β***_*g*_) and dispersion terms (*θ*_*g*_), which are estimated via GPU-accelerated variational inference. This structure allows for scalable inference while accounting for patient-level variability and sparse count data. **d.** The workflow for scDE analysis with DEVIL integrates variational inference and GPU for efficient estimation of model parameters, combined with robust covariance modelling to support high-throughput scDE expression analysis with control of false positives also in multi-patient scRNA-seq datasets.

The inputs that DEVIL requires are a matrix of raw gene expression counts ***Y*** of size *G* × *N* (with *y*_*g*,*n*_ being the count for gene *g* in cell *n* from one of *P* patients), and a design matrix ***X*** of size *N* × *F* encoding the experimental question of interest, such as cell type, treatment status, age, or any combination of categorical and continuous covariates. These ingredients are connected through a Negative Binomial GLM^18,19^, which models the mean expression of each gene as a log-linear function of the covariates, scaled by the size factors. For each gene *g*, counts are modelled as

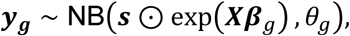

where *s* is a per-cell size factors vector that corrects for differences in sequencing depth^3,20,21^, ***β***_*g*_is the gene-specific coefficient vector quantifying the effect of each covariate on expression, *θ*_*g*_ is the gene-specific overdispersion, and ⊙ denotes the Hadamard (element-wise) product. DEVIL’s pipeline then proceeds in four stages (Fig.1d).

First, gene-specific expression coefficients, which quantify the effect of each covariate on expression, are inferred via a mean-field variational approximation^22–24^, an approach that, by design, recovers the classical iteratively reweighed least squares (IRLS) fixed point for Negative Binomial regression, while enabling efficient multi-GPU parallelization. Second, a gene-specific overdispersion parameter, capturing biological variability beyond Poisson noise, is estimated conditional on the inferred expression coefficients; DEVIL offers both a maximum-likelihood estimator (MLE) and a method-of-moments (MoM) alternative; the latter is algebraically simpler and GPU-compatible, offering consistent results with minimal loss of statistical efficiency. Third, the covariance of the expression coefficients is estimated to enable rigorous statistical testing: rather than relying on the variational posterior, which systematically underestimates uncertainty, DEVIL adapts its covariance estimator to the experimental design. In single-patient settings, the covariance is set to the inverse negative Hessian of the log-likelihood, the standard frequentist uncertainty estimate for a correctly specified GLM. In multi-patient settings, where cells from the same individual are correlated, DEVIL instead computes a clustered sandwich estimator, a method used in statistical analysis to provide robust standard error estimates when data is divided into clusters. This strategy, which treats patients as clusters, aggregates gradient contributions within each patient before estimating the empirical score covariance^16,17,25,26^. In particular, the covariance for the expression coefficients is computed as

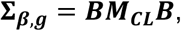

where ***B***, the inverse of the negative Hessian of the log-likelihood, acts as a ‘bread’ matrix, and ***M***_***CL***_ is the empirical covariance of the score vectors (‘meat’) computed across patient clusters. The key insight is that this estimator inflates standard errors to reflect the true number of statistically independent observations, i.e. the number of patients rather than the number of cells, without introducing any patient-specific parameters and keeping model complexity fixed regardless of cohort size, thereby avoiding the need for an extra number of parameters that arise in GLMs with mixed and fixed effects. Conceptually, this is equivalent to replacing the nominal cell count with a smaller effective sample size that reflects the degree of within-patient correlation. Fourth and finally, differential expression is assessed with Wald tests^27,28^, with p-values adjusted across genes using the Benjamini–Hochberg procedure^29^. Full model specification, variational update equations, and derivation of the sandwich estimator are provided in the Methods and in the Supplementary Information.

### Analyzing datasets with over 15 million cells and mixed experimental designs

We first evaluated the raw computational efficiency of DEVIL, demonstrating that its GPU-accelerated implementation achieves up to a 44.3x speedup with respect to the fastest scDE competitor and scales to datasets with up to 15 million cells. To benchmark performance, we compared runtime and memory usage of both the CPU and GPU versions of DEVIL against glmGamPoi^13^ and NEBULA^14^, leading methods for large-scale scDE analysis, across a range of dataset sizes. We based our benchmarking on a dataset from Chiou et al.^30^ containing 1.4 million cells, from which we derived 9 subsampled datasets ranging from 1,000 to 1,000,000 cells and from 100 to 5,000 genes (Methods). While all tools showed similar runtimes for small datasets (1,000 cells, 100 genes), performance diverged significantly at scale (Fig. 2a; Supplementary Table 3). At 1,000,000 cells and 1,000 genes, DEVIL’s GPU implementation completed analysis in 21.9 seconds, a 29.4x speedup over glmGamPoi (644 s) and 136x over NEBULA (2,981 s), using approximately 90% less memory than glmGamPoi, consistent with the latter’s reliance on large cell-by-gene matrix operations. The performance gap widened further with 5,000 genes, where DEVIL GPU completed analysis in ∼76 seconds versus glmGamPoi’s ∼57 minutes (3,458 seconds; 44.3x speedup for DEVIL) and NEBULA’s ∼200 minutes (12,221 seconds; 74.5x speedup for DEVIL). Importantly, to validate consistency between CPU and GPU implementations, we assessed that the two versions of DEVIL produced numerically identical parameters and p-values, and were statistically equivalent to glmGamPoi in single-patient settings (Supplementary Fig. 1); such agreement does not extend to multi-patient cases, where DEVIL’s clustered sandwich estimator leads to more conservative variance estimates and, consequently, different test results. For the same reason, discrepancies are also observed with respect to NEBULA.

**Figure 2.**
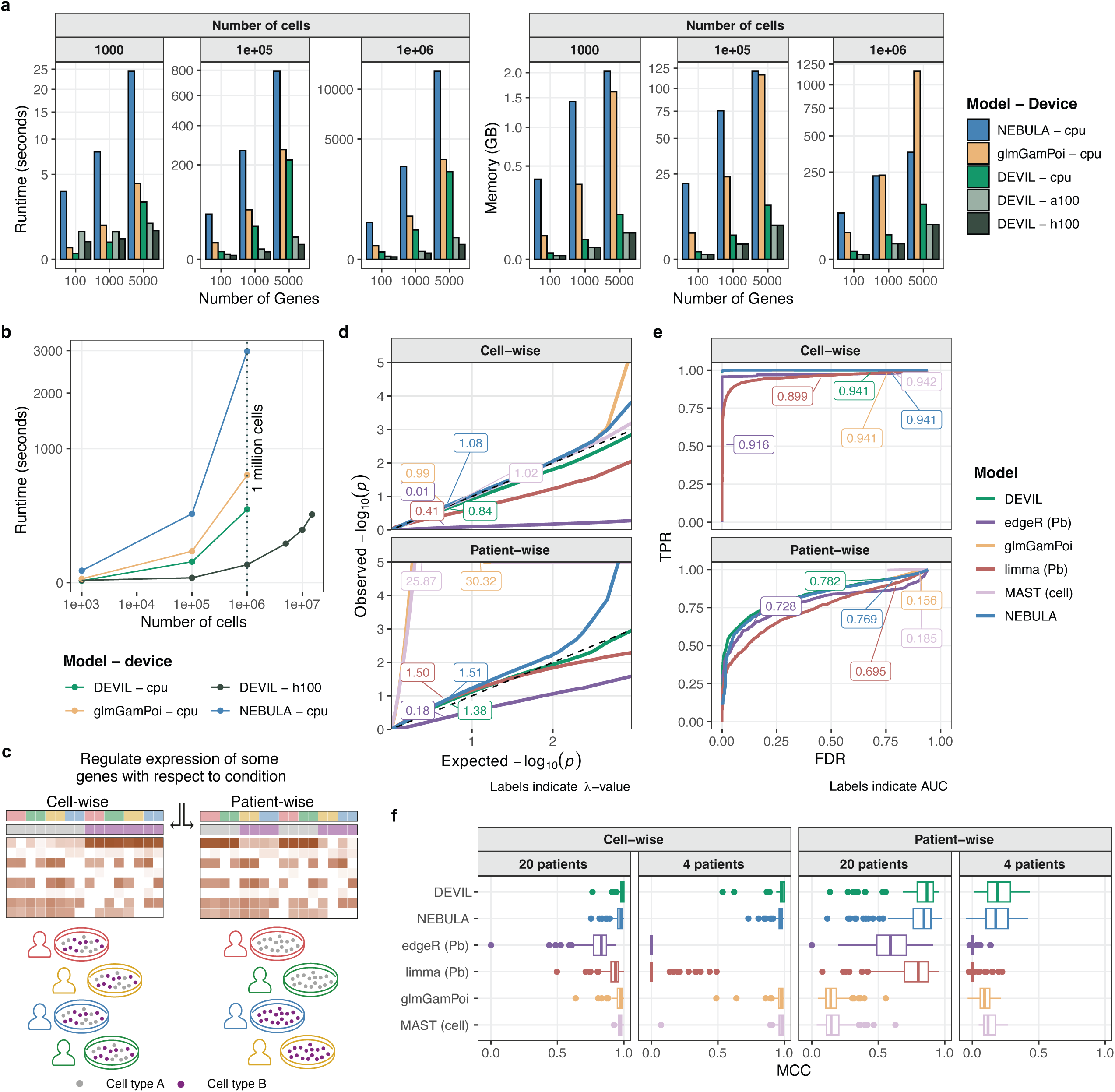
Performance and accuracy of DEVIL against state-of-the-art tools for scDE testing. **a.** Runtime and memory used by DEVIL as compared to glmGamPoi and NEBULA, for different input-dataset dimensions. Both CPU and GPU implementations of DEVIL are considered, as well as two distinct GPU architectures. DEVIL is both faster and more memory efficient. **b.** Absolute run-times with input dataset of 1000 genes and a growing number of cells. The GPU implementation enables the analysis of up to 15 million cells. **c.** Two contrasting experimental setups are considered for benchmarking multi-patient datasets. In the cell-wise design, both experimental conditions (e.g., cell types A and B) are represented within each patient, allowing within-patient comparisons that naturally control for inter-patient variability. In contrast, the patient-wise design assigns each condition to a distinct group of patients (i.e., each patient harbors only one condition) mimicking scenarios such as case-control studies or treatment comparisons across individuals. The latter introduces a complete confounding between condition and patient identity. **d, e.** QQ-plot (d) of expected vs. observed –log_10_ p-values for non-differentially expressed genes, with the inflation factor λ reported for each tool, and TPR/FDR curve (e) showing power of each tool for a nominal FDR threshold, with AUC values reported. DEVIL exhibits near-uniform distributions under the null hypothesis, while maintaining high power in detecting true DE genes. **f.** Performance evaluation using Matthews Correlation Coefficient (MCC) across varying numbers of patients (4 and 20) and experimental settings. DEVIL and NEBULA achieve top performances on both experimental settings. Boxes show the IQR with median indicated; whiskers extend to 1.5× IQR; points denote outliers.

To test limits of scalability, we applied DEVIL to datasets of up to 15 million cells and 1,000 genes. In this scenario, DEVIL’s GPU implementation demonstrated the ability to efficiently perform analyses previously impractical for existing tools (Fig. 2b). We further evaluated the strong scaling of DEVIL’s GPU implementation by varying batch size and number of GPUs. Runtime decreased approximately linearly with the number of GPUs, demonstrating efficient multi-GPU parallelization, and a batch size of 100 achieved near-optimal scaling while requiring less than 10 GB of GPU memory per device, enabling efficient execution on standard HPC systems (Supplementary Fig. 2). Results were consistent across an independent pancreas dataset^31^ (Supplementary Fig. 3, Supplementary Table 4).

Beyond scaling with respect to cell number, we further benchmarked computational performance as a function of patient counts and number of input features while keeping the total number of cells fixed, using simulated data to probe scenarios relevant to large cohort studies with high number of features to model (Methods). Across these additional scaling axes, we observed that increasing the number of patients had a pronounced effect on NEBULA’s runtime, reflecting the cost of subject-specific random effects, whereas DEVIL’s advantage over NEBULA remained stable as the number of features increased, indicating that performance gains are preserved as model complexity grows (Supplementary Fig. 4). Overall, these results establish DEVIL as an advancement in scRNA-seq analysis, offering performance improvements while maintaining optimal results.

### DEVIL controls false discoveries across mixed experimental designs

We next assessed DEVIL’s robustness to batch effects and mixed experimental designs through complementary simulation studies. In batch-effect experiments, we compared three modelling strategies: no batch correction, sandwich variance estimation alone, and sandwich estimation with explicit batch fixed effects (Methods). In small-scale regimes, sandwich estimation alone was sufficient to control false discoveries (median False Discovery Rate (FDR) below 0.05) while maintaining detection accuracy (median Matthews Correlation Coefficient (MCC)^32,33^ of 0.838), indicating that correcting standard errors is often adequate when batch effects primarily increase variance rather than induce systematic bias. In contrast, in large-scale regimes, batch effects became increasingly problematic when they introduced strong, systematic shifts in gene expression. While the sandwich estimator alone improved calibration compared to the plain model (reducing FDR from 0.944 to 0.293), explicit modelling of batch effects was required to fully restore both calibration and power. In this scenario, incorporating batch fixed effects achieved low FDR (<0.05) and high MCC (median of 0.837 compared to 0.692 obtained with sandwich estimation alone)(Supplementary Fig. 5, 6).

To benchmark DEVIL against the full landscape of scDE methods, we applied the simulation framework proposed by Lee et al.^34^ to four real scRNA-seq datasets^35–38^, comparing DEVIL against two pseudobulk-based methods (limma^5^ and edgeR^4^) and three single-cell-based methods (glmGamPoi^13^, NEBULA^14^ and MAST^39^). This setup tested tools both within and outside their intended use cases to evaluate their robustness. The framework considered two critical scenarios: *cell-wise*, where each patient contains cells from all conditions (e.g., cell-types comparison), and *patient-wise*, where each patient belongs to a single condition (e.g., case-control), creating perfect confounding between patient and treatment (Fig. 2c). In both settings, pseudobulk counts are obtained by aggregating cells within each (patient, condition) group, ensuring DE is evaluated at the correct replicate level, whereas single-cell methods operate directly on individual cells. Performance was assessed via quantile-quantile (QQ) plots of p-values under the null, expecting a uniform distribution, by analyzing true positive rate (TPR) versus FDR curves (iCOBRA^40^; Fig. 2d,e), and by measuring MCC across varying patient numbers and DE fractions (Fig. 2f).

In both scenarios, DEVIL achieved robust results thanks to its sandwich covariance estimator, which enables it to handle the patient-wise scenario correctly, accounting for within-patient correlation. Other tools revealed limitations arising from their underlying assumptions. For instance, limma and edgeR showed type II errors in *cell-wise* settings (i.e., genes truly differentially expressed were not identified) because pseudobulk discards within-patient, between-cell variability information that is crucial for detecting subtle expression differences. Conversely, glmGamPoi and MAST exhibited type I errors in *patient-wise* scenarios (i.e., genes were misleadingly identified as differentially expressed), due to their treating of cells as independent observational units, which leads to systematic underestimation of coefficient uncertainty when cells are correlated within patients, resulting in severely inflated test statistics and p-values (Fig. 2d,e). NEBULA performed well in both scenarios due to its subject-specific random effects, but at considerably higher computational cost. For example, taking as example the Suo et al.^35^ dataset with 20 patients (*n=90* simulations per setting), in cell-wise settings it reached a median MCC of 0.990 ([0.943, 1], 5–95%), comparable to NEBULA (0.978 [0.881, 1]), glmGamPoi (0.976 [0.881, 1.000]), and MAST (0.973 [0.941, 1.000]), and outperforming pseudobulk methods (limma: 0.937 [0.773, 0.979]; edgeR: 0.830 [0.598, 0.911]), which discard within-patient between-cell variability and consequently lose power for detecting subtle expression differences. In the more challenging patient-wise scenario, DEVIL achieved a median MCC of 0.866 [0.370, 0.947], modestly outperforming NEBULA (0.844 [0.358, 0.941]) and limma (0.801 [0.377, 0.931]), and substantially outperforming glmGamPoi (0.142 [0.068, 0.335]) and MAST (0.145 [0.071, 0.360]), which treat cells as independent observations and therefore severely underestimate coefficient uncertainty when cells are correlated within patients, leading to inflated test statistics. This test clearly shows that our approach can mitigate patient-specific noise, i.e. systematic expression differences from individual patient characteristics rather than experimental conditions (Fig.2f). From a computational cost perspective, DEVIL achieved these results while being faster than scDE competitors (median speedup: 4.433 [3.581, 5.707] over NEBULA; 2.363 [1.889, 2.934] over glmGamPoi; 10.813 [9.125, 12.959] over MAST). These patterns were consistent across ∼1,500 datasets from four independent cohorts^35–38^ and additional competing methods (Supplementary Figs. 7-11; Supplementary Tables 5-9), where pseudobulk approaches consistently underperformed in cell-wise settings due to loss of within-patient information, while single-cell methods treating cells as independent observations (e.g. glmGamPoi and MAST) frequently exhibited inflated p-values. Within DEVIL, MLE and MoM overdispersion estimators yielded indistinguishable accuracy across all simulations, with MoM offering consistent runtime improvements (Supplementary Fig.12). Notably, NEBULA was generally comparable to DEVIL, highlighting the flexibility of the mixed-effect implementation. However, DEVIL offered the best trade-off across statistical performance and computational efficiency, establishing it as a robust and scalable framework for DE analysis in single– and multi-patient scRNA-seq studies.

### Correcting for multi-patient variability enhances reliable and precise cell-type marker detection

A common downstream task in scRNA-seq analysis is the identification of cluster-specific marker genes for cell-type annotation, a step highly sensitive to the quality and specificity of DE calls, which can vary substantially across methods. To assess whether DEVIL’s statistical framework translates into more accurate and biologically grounded cell-type annotation, we evaluated its performance across datasets with independent or experimentally validated labels, trying to mitigate double-dipping and label circularity, using the three best-performing tools from our earlier benchmarks: DEVIL, NEBULA, and glmGamPoi.

We first used the multimodal PBMC dataset of Hao et al.^41^, where CITE-seq–derived protein measurements provide robust, modality-independent cell-type labels, reducing reliance on RNA-only clustering and limiting label circularity. Cells were clustered with Seurat^42^ (Fig. 3a), and DE genes were computed per cluster using each method. The top N DE genes per cluster, selected by log₂ FC ≥ 1.0 and p-value threshold, were passed to scMayoMap^43^ for annotation (Fig. 3b-d). Accuracy was evaluated across a systematic grid of marker set sizes (5 to 1,000 genes) and p-value thresholds (Methods). In this setting, DEVIL achieved the highest mean annotation accuracy at 25 markers per cluster (50.7 ± 0.7%), followed by NEBULA at 5 markers (50.0 ± 3.1%) and glmGamPoi at 25 markers (47.6 ± 0.03%). Peak accuracies were 53.5% for DEVIL and 51.6% for NEBULA (both at 5 markers, p = 0.01), and 47.6% for glmGamPoi at 25 markers with p = 0.01. For small, high-confidence marker sets (5–10 genes), DEVIL and NEBULA performed comparably, reflecting their shared ability to extract specific and discriminative markers. However, for sets of 25 genes or more, DEVIL consistently outperformed both across all thresholds. As the number of markers increased further, accuracy for all methods gradually declined and converged toward ∼40% for DEVIL and ∼35% for the others (Fig. 3e), reflecting the increased ambiguity introduced by broader, less specific gene lists. In general, DEVIL was more conservative than glmGamPoi in the total number of DE genes called per cluster, while NEBULA showed markedly inconsistent behavior, for example, detecting far fewer DE genes in some clusters (cluster 12: 469 vs. 1,089 for DEVIL) and many more in others (cluster 13: 3,201 vs. 791) (Fig. 3f), likely reflecting instability in variance component estimation. Additionally, in terms of runtime, DEVIL was on average 3.5x faster than NEBULA (Fig. 3g), confirming its scalability advantage. Results were consistent across three additional annotated datasets (Lee et al.^44^, Guilliams et al.^45^ and MacParland et al.^46^; Supplementary Fig.13-15), with DEVIL and NEBULA alternating at the top but DEVIL consistently offering the best trade-off between accuracy and computational efficiency.

**Figure 3.**
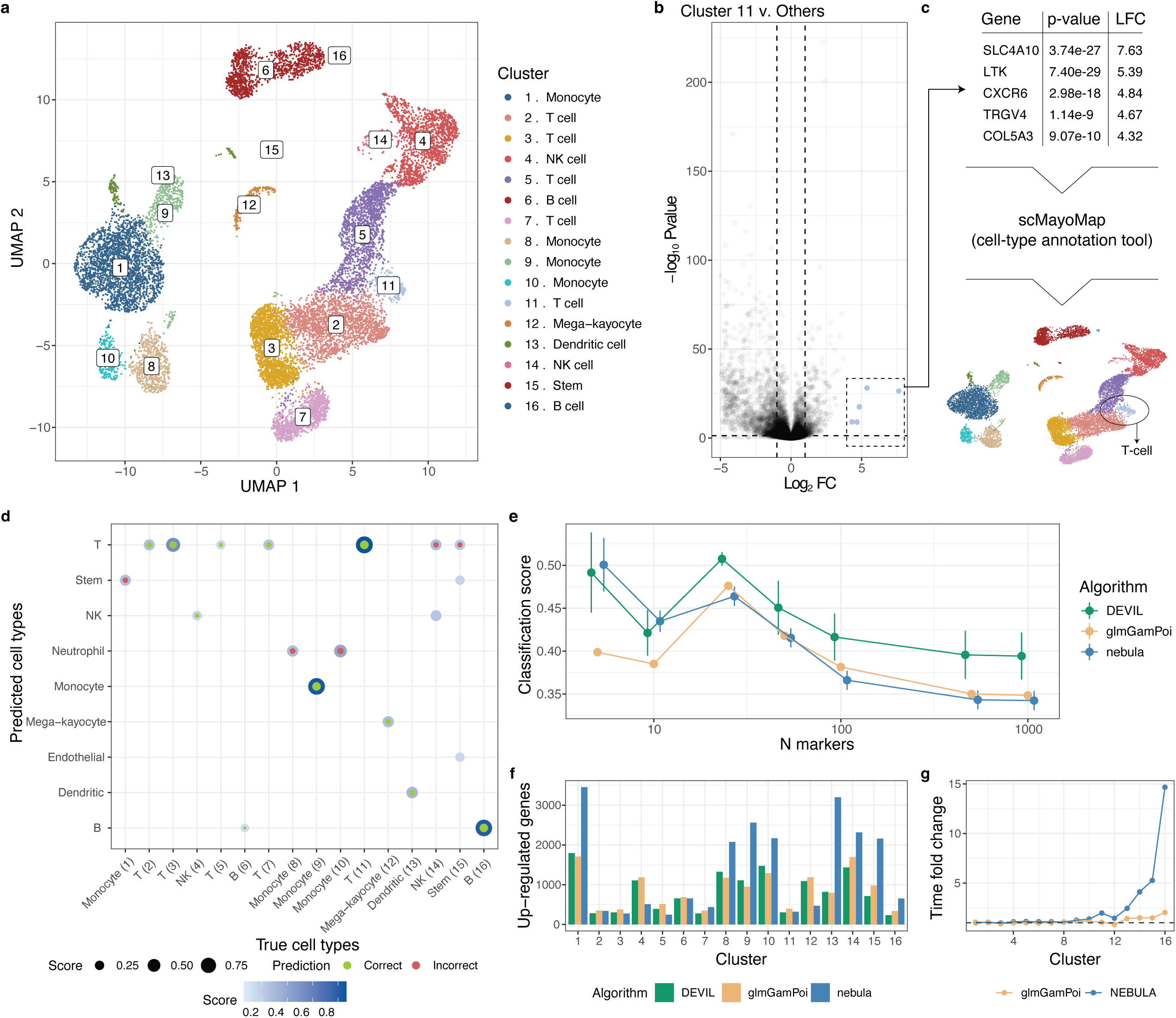
Identification of cell types from scDE markers of PBMC cells of multiple patients. **a.** UMAP visualization of cell populations published by Hao et al.^41^. Sixteen transcriptionally distinct clusters were identified and annotated using ground truth cell type labels, including T cells, B cells, monocytes, and additional immune and stromal populations. **b, c.** Differential expression in cluster 11. A volcano plot displays gene-level LFC and statistical significance for cluster 11 versus all other clusters. Five marker genes (highlighted in light blue) exceed both expression magnitude and significance thresholds, indicating strong cluster specificity. The identified marker genes are used for automated cell type annotation with scMayoMap, indicating that cluster 11 contains T-cells, consistent with its transcriptomics profile and spatial location on the UMAP in panel (a). **d, e.** Predicted versus true cell types for automated annotation using scMayoMap. The classification matrix (d) compares predicted (y-axis) and true (x-axis) cell types. Circle size and color denote classification confidence, with green indicating correct and red indicating incorrect predictions. The classification accuracy (e) is compared across DEVIL, glmGamPoi, and NEBULA as a function of marker set size. Points show mean accuracy, and error bars denote ±1 s.d. across *n=4* p-value thresholds to reflect parameter sensitivity of the analysis. **f.** The number of differentially expressed genes identified per cluster varies by method, highlighting differences in stringency and sensitivity of each tool tested. **g.** Relative runtime (normalized to DEVIL) across methods, combined with precision estimates in panel (e), suggests a trade-off between speed and precision. glmGamPoi is fast but less precise than DEVIL, whereas NEBULA is precise but shows increased computational cost in larger clusters.

To complement the multimodal analysis and assess true marker-recovery capability against an unambiguous ground truth, we next analyzed the Liu dataset^47^, in which 10 immune cell populations were isolated through antibody-based positive and negative selection, uniquely barcoded, and pooled prior to sequencing. Because each population is experimentally purified, this dataset provides true, validated cell-type labels entirely independent of computational clustering. Hence, DE genes were computed directly from known labels without clustering, and each method’s ranked gene list was evaluated against curated marker sets using ROC curves and AUC (Methods; Fig. 4a).

**Figure 4.**
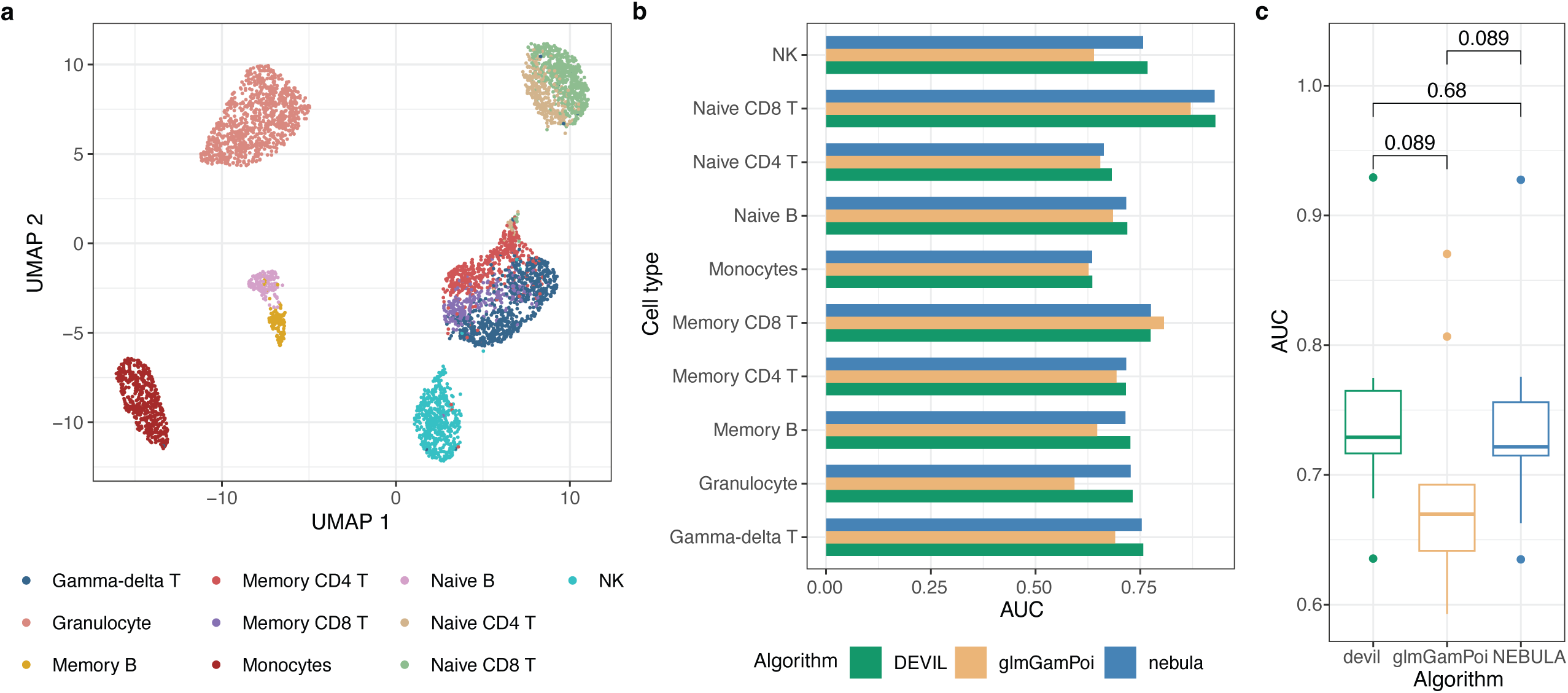
Marker gene recovery in purified immune cell populations. **a.** UMAP embedding of the Liu dataset^47^ comprising 10 immune cell populations isolated by antibody-based positive and negative selection and uniquely barcoded prior to pooling and sequencing. Colors indicate experimentally validated cell-type identities, which serve as ground truth labels. **b.** Area under the ROC curve (AUC) for marker gene recovery for each cell type, comparing DEVIL, glmGamPoi, and NEBULA. For each method, ROC curves were constructed by ranking genes by differential expression statistics and evaluating against curated binary marker annotations. **c.** Distribution of AUC values across all cell types for each method. Reported p-values denote pairwise comparisons between methods (Wilcoxon signed-rank test). Boxes show the IQR with median indicated; whiskers extend to 1.5× IQR; points denote outliers.

At the level of individual cell-types, glmGamPoi surpassed the other methods only for memory CD8+ T cells (AUC = 0.806 vs. 0.775 for DEVIL and NEBULA, Fig. 4b). For all remaining populations, DEVIL and NEBULA achieved superior and more consistent performance, likely reflecting their ability to correctly model within-patient correlation structure and thereby produce more stable and biologically consistent marker rankings. In fact, DEVIL achieved the highest mean AUC (0.744), followed closely by NEBULA (0.738), both outperforming glmGamPoi (0.690) (Fig. 4c). Overall, DEVIL delivered the strongest aggregate performance with lower variability across cell types, confirming its robustness in recovering validated DE markers.

### Distilling functionally relevant transcriptional signatures in ageing muscle

We applied DEVIL, glmGamPoi, and NEBULA to single-nucleus RNA sequencing (snRNA-seq) data from human limb skeletal muscle of 19 individuals spanning a broad age range^48^. To characterize ageing-associated transcriptional changes, we fitted a linear interaction model between age group (young vs. old) and myofiber subtype (Type I vs. Type II), the two most abundant myonuclei populations. Dimensionality reduction revealed clear separation between subtypes, while age-dependent differences were subtler and partially overlapping (Fig. 5a,b).

**Figure 5.**
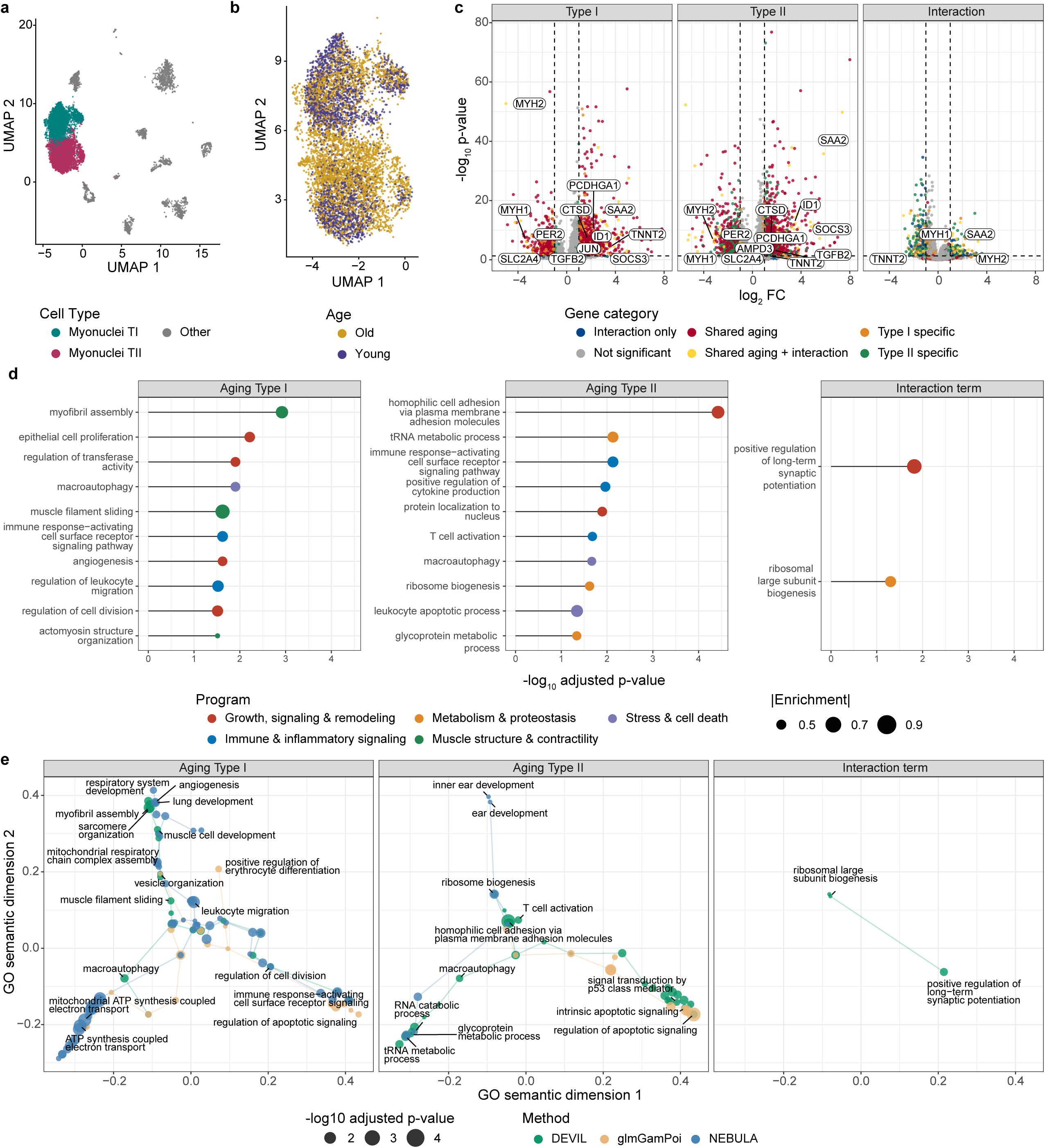
Ageing-associated transcriptional programs in human skeletal muscle myonuclei. **a, b.** UMAP embedding of the snRNA-seq dataset from human limb skeletal muscle, colored by myonuclei subtype (Type I, Type II, other; a) or age group (young vs. old; b). **c.** Volcano plots showing differential expression results obtained with DEVIL for ageing contrasts in Type I myonuclei, Type II myonuclei, and the age-by-myofiber subtype interaction. Genes are colored by category and selected biologically relevant markers are annotated. **d.** Top enriched Gene Ontology biological process terms identified by DEVIL for each contrast. Dot position indicates significance, dot size reflects enrichment magnitude, and colors denote broad biological program categories. **e.** Two-dimensional GO semantic similarity maps summarizing enriched pathways across contrasts and methods. Each point represents a GO term positioned by semantic similarity, with point size indicating statistical significance. Adjusted P values were obtained from Gene Set Enrichment Analysis (GSEA; permutation-based test with 50,000 permutations) and corrected for multiple comparisons using the Benjamini–Hochberg (BH) procedure.

The original study highlighted a panel of biologically relevant marker genes spanning muscle contractile machinery (MYH1, MYH2), sarcomeric structure (TNNT1, TNNT2), adhesion and inflammatory response (SAA1, SAA2, PHLDB2), and major signaling pathways (PDE4B, JUN, SOCS3). Their expression patterns were preserved in our analysis, with MYH7B enriched in Type I myonuclei, MYH2 in Type II, and JUN upregulated in aged cells. These expected results were summarized by DEVIL: for example, SAA2 was strongly enriched in aged myocytes, particularly Type II, while MYH1 and MYH2 exhibited decreased expression with age (Fig. 5c; Fig.6a). All three methods recapitulated established single-gene signatures from the original study, confirming that core ageing biology was captured at the single-gene level across all tools (Supplementary Fig. 16). To compare pathway-level interpretations, we performed Gene Set Enrichment Analysis (GSEA) and embedded enriched Gene Ontology (GO) terms into a two-dimensional semantic similarity space to analyse the biological axes recovered by each method (Methods).

In Type I myonuclei, DEVIL recovered a highly coherent muscle-specific ageing program: enriched processes spanned sarcomeric remodeling (myofibril assembly, muscle filament sliding, actomyosin organization), cell-cycle and turnover regulation, and macroautophagy (Fig. 5d), pathways that align with established hallmarks of slow-twitch fibre ageing, including structural degeneration and proteostatic surveillance^49,50^. In semantic space, GO terms formed three interpretable axes: 1) mitochondrial and ATP synthesis pathways, 2) immune and inflammatory signaling, and 3) tissue organization and developmental programs. NEBULA also detected mitochondrial and bioenergetic pathways, often with strong statistical signal, but introduced less tissue-specific developmental programs, including terms related to lung and respiratory morphogenesis, that are difficult to reconcile with skeletal muscle biology. In contrast, glmGamPoi primarily recovered apoptotic and cell-cycle processes, failing to capture the structural and tissue-organization programs that are central to Type I fibre ageing (Fig. 5e; Supplementary Fig. 17, 18).

In Type II myonuclei, DEVIL again identified a multifaceted and biologically consistent response characterized by translational remodeling (ribosome biogenesis^51^, tRNA metabolism), immune activation (T-cell activation, cytokine production), macroautophagy, and homophilic cell adhesion^49^, signatures consistent with the preferential atrophy of fast-twitch fibres in ageing and with the proteostatic dysregulation and altered intercellular communication that characterize sarcopenic muscle^49^. Once again, glmGamPoi was dominated by apoptotic signaling, while NEBULA introduced off-target development terms (e.g., inner ear formation), suggesting reduced tissue specificity. Interestingly, only DEVIL detected significant GO terms specific to the age-by-cell-type interaction, including ribosomal large subunit biogenesis and positive regulation of long-term synaptic potentiation, processes consistent with loss of neuromuscular junction integrity and muscle atrophy^52,53^ (Fig. 5d,e; Supplementary Fig. 17, 18).

Across the two major cell-types, Type I and Type II, we performed additional analyses to assess the stability, specificity, and biological fidelity of the enriched pathways detected by each method. In Type I myocytes, DEVIL recovered the highest number of muscle-relevant biological processes, with NEBULA ranking second and glmGamPoi failing to identify any muscle-specific pathways (Fig. 6b). In Type II myocytes, all methods predominantly enriched more generic processes (translational, metabolic, and stress-response pathways), with no clear advantage in muscle specificity for any single tool.

**Figure 6.**
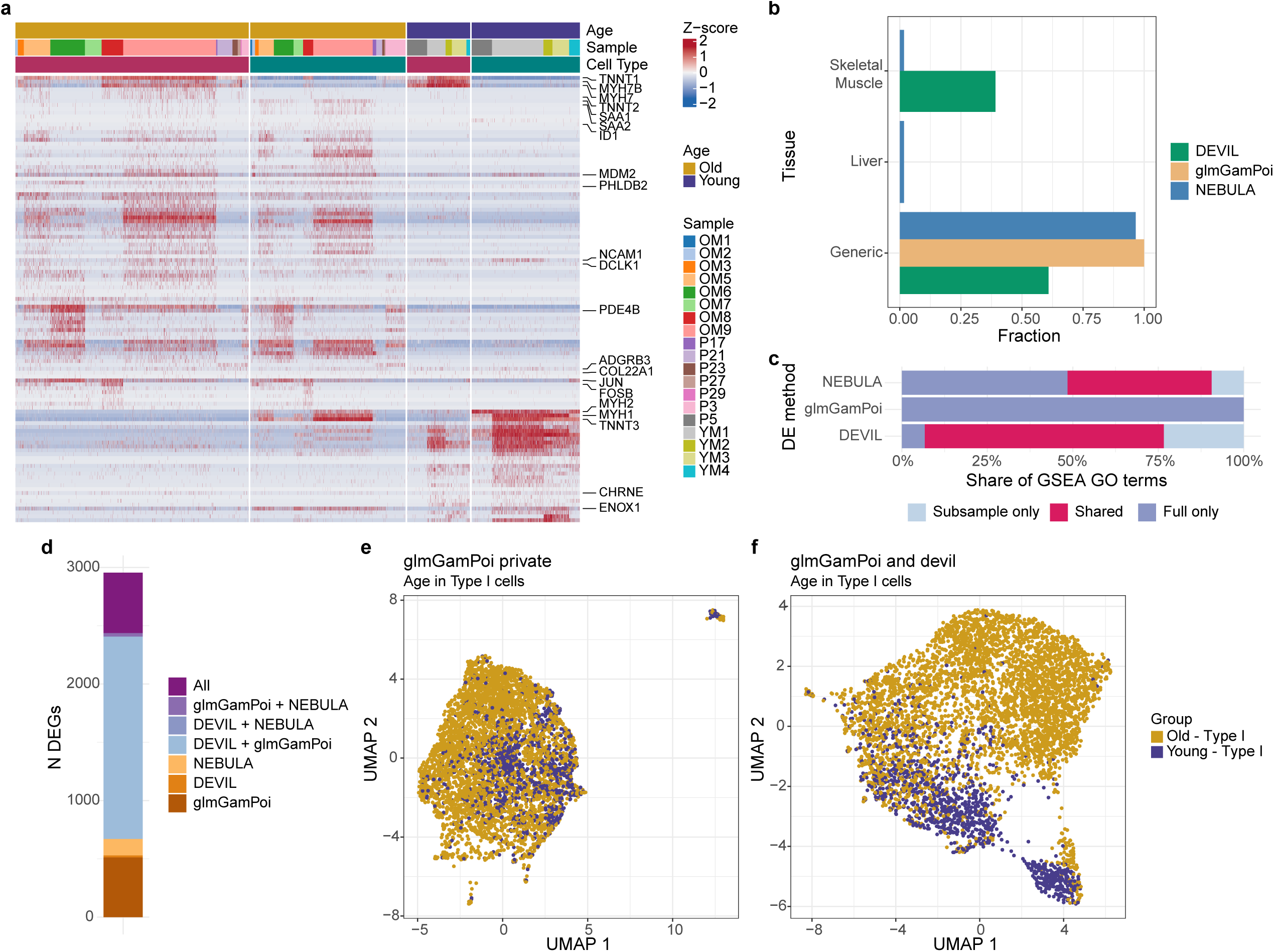
Robustness and specificity of ageing-associated signals. **a.** Heatmap of z-score–normalized expression for the top ageing-associated differentially expressed genes identified by DEVIL, with cells annotated by age, sample, and myofiber subtype. **b.** Fraction of enriched Gene Ontology biological process terms classified as skeletal muscle–specific, liver-related, or generic for ageing in Type I myonuclei, across methods. **c.** Overlap of enriched GO terms between full and subsampled datasets for ageing in Type I myonuclei, showing terms detected only in the full dataset, only in subsampled data, or shared between the two. **d.** Number of differentially expressed genes uniquely identified by each method or shared across methods for ageing in Type I myonuclei. **e, f.** UMAP embeddings constructed using top 100 DE genes identified exclusively by glmGamPoi (e) or shared between glmGamPoi and DEVIL (f), with cells colored by age group.

To assess robustness to data imbalance, we repeated all analyses on subsampled datasets equalizing cell numbers across age groups and myofiber types. While the number of enriched pathways varied, DEVIL consistently showed the largest overlap between subsampled and full data GO term sets, followed by NEBULA, with glmGamPoi showing dramatically reduced reproducibility (Fig. 6c). This was true not only for Type I cells, but also in Type II, highlighting the methodological stability of DEVIL in realistic scenarios of uneven cell representation (Supplementary Fig. 19).

Finally, to investigate whether DEVIL’s conservatism relative to glmGamPoi reflects genuine signal filtering rather than power loss, we compared low-dimensional embeddings constructed from glmGamPoi-private DE genes versus genes shared between glmGamPoi and DEVIL (Fig. 6d). Embeddings based on glmGamPoi-only genes showed modest age-related separation, whereas shared genes produced a substantially cleaner and crisper ageing gradient, consistent across both myofiber subtypes (Fig. 6e,f; Supplementary Fig. 20). This indicates that DEVIL selectively retains the biologically meaningful portion of glmGamPoi’s signal while discarding possibly noisy detections.

Together, these analyses show that DEVIL provides the most biologically focused and muscle-relevant interpretation of ageing across both myofiber subtypes. Compared to alternative tools, DEVIL consistently reduces noise, avoids tissue-irrelevant developmental enrichments, and prioritizes pathways that align with established mechanisms of skeletal muscle ageing. Overall, these results position DEVIL as a robust framework for extracting coherent biological programs from complex, heterogeneous single-cell datasets.

## Discussion

Single-cell RNA sequencing (scRNA-seq) has revolutionized our ability to resolve cellular heterogeneity; however, statistically robust differential expression (DE) analysis across individuals remains a fundamental challenge. Traditional DE frameworks, designed for bulk data, assume independence across samples, a premise that is violated in scRNA-seq, where thousands of correlated observations arise from a single biological replicate. Ignoring these dependencies can inflate false discovery rates and undermine population-level inference.

These limitations are overcome by DEVIL, which combines a GLM with clustered sandwich variance estimation, enabling scalable and rigorous scDE testing while preserving cell-level resolution. Unlike pseudobulk methods, which obscure intra-sample variation, or mixed-effect models, which scale poorly with patient number, DEVIL accurately quantifies uncertainty by treating patients as the unit of replication and modelling intra-patient correlations without increasing model complexity. Moreover, DEVIL is flexible enough to support both patient-wise (e.g., case-control) and cell-wise (e.g., perturbation) designs through a unified statistical framework. By adjusting for subject-level correlation without requiring subject-specific parameters, DEVIL maintains statistical power while keeping model complexity fixed regardless of cohort size.

Across a wide range of datasets, including simulations, immune profiling, and large-scale tissue atlases, DEVIL matches or outperforms established GLM-based and pseudobulk methods across all evaluated settings. Moreover, DEVIL is, to our knowledge, one of the few cell-level methods capable of efficiently analyzing datasets exceeding 15 million cells and thousands of genes. We note that pseudobulk-based approaches^3–5,54^ can also process nominally large cell counts rapidly, but their computational cost scales with the number of patients rather than the number of cells, conferring an inherent scalability advantage that comes at the cost of sacrificing within-patient cell-to-cell variability. This makes DEVIL a practical solution for modern scRNA-seq studies involving millions of cells and requiring both cell-level resolution and population-scale inference, such as large-scale atlas efforts^30,48,55,56^ or drug screening assays^57–59^.

Despite DEVIL’s methodological and computational advances over the state of the art, challenges remain, especially at the methodological level. The current model assumes gene-specific but patient-invariant overdispersion, which may limit its ability to capture individual-specific regulatory effects. As with all model-based approaches, performance may degrade in the presence of severe unmodeled confounding, batch effects, or sparsity. In addition, as in most current scRNA-seq differential expression frameworks, our analyses assume that cell-level annotations (e.g. cell types) are known and treated as fixed. While common practice, errors in cell labelling, particularly when labels are derived from clustering, can propagate into downstream DE testing and inflate significance. Addressing label uncertainty and jointly accounting for clustering and differential expression remains an open methodological challenge and lies beyond the scope of the present work but could be integrated in future extensions of DEVIL through uncertainty-aware cell representations. Future extensions could introduce hierarchical priors, interaction terms, or nonlinearities to improve flexibility, while maintaining computational efficiency. Similarly, the principles of DEVIL could be adapted to model other types of single-cell assays, such as those probing chromatin accessibility^60–62^.

Moreover, as single-cell technologies evolve toward longitudinal, spatial, and multi-omics profiling, future iterations of DEVIL should incorporate temporal structure, multi-modal integration, and phylogeny-aware priors, especially in complex contexts such as cancer. Our results demonstrate that principled, scalable inference is achievable without sacrificing resolution, providing a foundation for population-scale single-cell studies that demand both statistical rigor and computational tractability.

## Methods

### Overview of DEVIL model

#### Statistical framework

Raw single-cell RNA-seq count data are modelled by DEVIL using a Negative Binomial (NB)^18,19^ GLM, suitable for both single– and multi-patient experimental designs. The framework requires three inputs: (i) a raw RNA count matrix ***Y*** of size *G* × *N*, where *G* is the number of genes and *N* is the number of cells, with *y*_*g*,*n*_ being the counts for gene *g* in cell *n*; (ii) a design matrix ***X*** of size *N* × *F*, where *F* is the number of features, encoding experimental covariates, which may be both categorical and continuous (e.g., cell type, treatment status, age); and, for multi-patient data, (iii) a patient assignment vector ***p*** of size *N*, mapping each cell to its source individual. For each gene *g*, counts are modelled as:

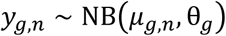

where *θ*_*g*_ is the gene-specific overdispersion and μ_*g*,*n*_ is the mean expression, defined as the *n* – th element of the vector

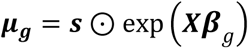

where ***β***_*g*_ is the gene-specific coefficient vector quantifying the effect of each covariate on expression. Per-cell size factors vector *s* of size *N*, correcting for differences in sequencing depth, is computed prior to fitting using standard normalization schemes (e.g., CPM, geometric-mean scaling, or single-cell–specific normalizations^20,21^) and incorporated into the expected mean expression through Hadamard ⊙ (element-wise) product; for notation brevity, all equations below assume *s* = 1, i.e. μ_*g*_ = exp (***X******β***_*g*_), without loss of generality. This parameterization implies a variance of μ + μ^2^/*θ*, capturing both Poisson sampling noise and biological overdispersion. Mathematical notation is summarized in Table 1.

**Table 1.** Mathematical notation of the DEVIL framework.

| Parameter | Symbol | Type | Dimension |
| --- | --- | --- | --- |
| Input matrix | $\mathbf{Y}$ | Matrix | $\mathbb{R}^{G \times N}$ |
| Design matrix | $\mathbf{X}$ | " | $\mathbb{R}^{N \times F}$ |
| Expression covariance | $\Sigma_{\beta}$ | " | $\mathbb{R}^{F \times F}$ |
| Bread matrix | $\mathbf{B}$ | " | $\mathbb{R}^{F \times F}$ |
| Meat matrix | $\mathbf{M}_{\text{CL}}$ | " | $\mathbb{R}^{F \times F}$ |
| Cell-to-patient assignment | $\mathbf{p}$ | Vector | $\mathbb{R}^N$ |
| Mean expression | $\mu_g$ | " | $\mathbb{R}^N$ |
| Scaling factor | $s$ | " | $\mathbb{R}^N$ |
| Expression coefficients | $\beta_g$ | " | $\mathbb{R}^F$ |
| Feature/Covariate vector | $\mathbf{x}_n$ | " | $\mathbb{R}^F$ |
| Lambda vector | $\lambda_g$ | " | $\mathbb{R}^N$ |
| Omega vector | $\mathbf{w}_g$ | " | $\mathbb{R}^N$ |
| Overdispersion | $\theta_g$ | Scalar | $\mathbb{R}$ |

Equivalently, the NB distribution arises as a Gamma-Poisson mixture:

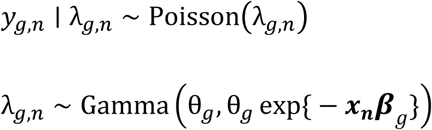

where λ_*g*,*n*_ is a latent, cell-specific Poisson rate. Intuitively, λ_*g*,*n*_ can be thought of as the unobserved true expression of gene g in cell n, and the Gamma prior encodes not only the expectation from the GLM but also, through the overdispersion θ_*g*_, how much individual cells are allowed to deviate from it.

#### Expression coefficient inference

For each gene, DEVIL estimates ***β***_*g*_using a mean-field variational approximation to the posterior, exploiting a Poisson-Gamma mixture representation of the NB distribution. This approximation recovers the classical iteratively reweighted least squares (IRLS) fixed point for NB regression, ensuring statistical equivalence to standard GLM fitting while admitting efficient multi-GPU parallelization. The update scheme iterates between estimating the auxiliary vector λ_**g**_ and updating the coefficient mean ***β***_*g*_ and covariance ***Σ***_***β***_. Given the current estimate of ***β***_*g*_ and ***Σ***_***β***_ one first computes:

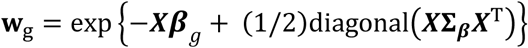

and then, the full update scheme becomes:

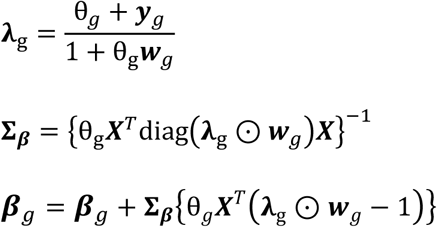

Above, we used the diag(⋅) and diagonal(⋅) notation, where diag(⋅) takes a vector and puts it on the diagonal of a zero matrix while diagonal(⋅) does the opposite, extracting the diagonal entries of a matrix as a vector. Full derivation of these updates and pseudocode are provided in Supplementary Information.

#### Overdispersion inference

Overdispersion *θ*_*g*_ is estimated conditional on ***β***_*g*_ using either an MLE or a MoM estimator. In the ML approach, *θ*_*g*_ is obtained by maximizing the negative binomial log-likelihood conditional on the estimated means, a two-step procedure that is widely used in bulk and single-cell RNA-seq^3,4,13,63^. While the mean and dispersion parameters are not fully separable in the negative binomial model, this approximation is well justified in practice, as dispersion primarily influences the variance structure and can be consistently estimated conditional on reasonable mean estimates. However, this method requires iterative optimization involving digamma and trigamma functions that cannot be efficiently offloaded to GPUs. As an efficient alternative, the MoM estimator equates the empirical variance of residuals *r*_*g*,*n*_ = *y*_*g*,*n*_ − μ_*g*,*n*_ to the model-implied variance, yielding a closed-form estimate:

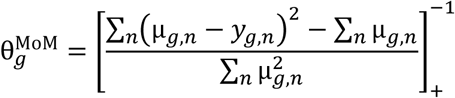

Where [⋅]_+_ denotes truncation to positive values. The MoM requires only elementwise operations and reductions. Hence, while less efficient than MLE in small samples, this estimator avoids iterative optimization and is well suited for large-scale analyses and GPU acceleration (Supplementary Information).

#### Covariance estimation and hypothesis testing

Accurate estimation of ***Σ***_***β***_ is crucial for differential expression testing. The variational posterior of ***Σ***_***β***_ is not directly used for hypothesis testing, as it typically underestimates uncertainty due to the mean-field assumption and, more importantly, does not incorporate the effect of *θ*_*g*_ on variance estimation. Even in the single-patient case, failing to account for the correct overdispersion leads to underestimated standard errors and inflated test statistics. Hence, DEVIL estimates ***Σ***_***β***_ using one of two approaches depending on experimental design. For single-patient data, ***Σ***_***β***_ is set to the inverse negative Hessian of the log-likelihood:

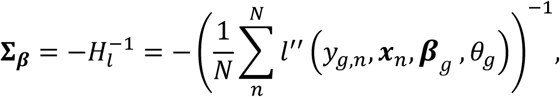

where *l*^′′^is the second derivative of the log-likelihood with respect to ***β***. For multi-patient data, DEVIL computes a clustered sandwich estimator^16,17^:

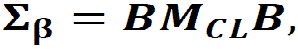

where ***B*** is the inverse negative Hessian (bread) and ***M***_***CL***_ is the empirical covariance of patient-level score sums (meat):

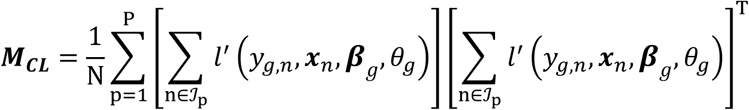

where *l*^′^ is the gradient of the log-likelihood with respect to ***β***, and ℐ_p_ denotes the set of cell indices for patient p. By aggregating gradient contributions within each patient before forming the outer product, this estimator accounts for within-patient correlations without introducing patient-specific parameters, in contrast to fixed– or mixed-effects models, which both require additional parameters. The resulting inflation of ***M***_***CL***_ relative to the standard (unclustered) meat can be interpreted as a reduction to an effective sample size *n*_eff_ < *N*, reflecting the true degree of independent information in the data (Supplementary Information). Hence, although DEVIL does not fit an explicit random-effects model, the clustered sandwich estimator introduces an empirical form of shrinkage across patients. This mirrors the core behavior of mixed models, enabling robust inference over hierarchical structures while avoiding the complexity of fully specified random-effects frameworks.

Differential expression is assessed via Wald tests for contrasts of the form *H*_0_: *c**β*** = 0, using the test statistic 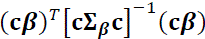, which follows a chi-squared distribution asymptotically. After this, *p*-values are adjusted using the Benjamini–Hochberg procedure to control the false discovery rate across genes. While DEVIL uses variational inference to estimate ***β*** efficiently, hypothesis testing is performed using Wald statistics with robust standard errors. This hybrid approach combines the scalability of VI and the robustness of classical uncertainty quantification.

### scRNAseq datasets

All the datasets used were obtained from publicly available databases and are summarized in Supplementary Table S2.

### Computational scalability benchmarking

#### Data preparation

Scalability was benchmarked using two datasets. The macaque brain dataset (Chiou et al.^30^, 1,437,348 cells, 15,562 genes) was used for large-scale evaluation. The human pancreas dataset (Baron et al.^31^, 8,569 cells, 20,125 genes) provided an independent replication at smaller scale. For the macaque brain dataset, the two most abundant cell types were retained; cells with fewer than 100 total RNA counts and genes expressed in fewer than 100 cells were removed. A two-variable design matrix encoding cell type was constructed. For the pancreas dataset, alpha and beta cells were selected; genes expressed in fewer than 100 cells were removed and a cell-type design matrix was used.

#### Scaling configurations

For the macaque brain dataset, we evaluated all combinations of {100, 1,000, 5,000} genes and {1,000, 100,000, 1,000,000} cells. For the pancreas dataset, combinations of {100, 1,000, 10,000} genes and {500, 1,000, 4,000} cells were tested. For each configuration, cells were ranked by total RNA count and genes by cumulative expression across retained cells, then subsampled accordingly. Runtime and peak memory were measured using the bench R package^64^ for DEVIL (CPU and GPU), glmGamPoi, and NEBULA. Scalability to 15 million cells was assessed using the macaque brain dataset with replication to match the target size. Multi-GPU scaling was evaluated by varying batch size ({100, 200, 500,1000} genes per batch) and GPU count ({1, 2, 4,8} devices).

#### Scaling across patients and features

Single-cell RNA-seq count data were simulated using a Splatter-based framework^65^, generating datasets of 10,000 cells and 1,000 genes across 8 to 50 patients (cells distributed approximately evenly). Experimental designs included 1 to 5 binary covariates, with Gaussian noise added to a subset to emulate continuous predictors. Ten independent replicates were generated per configuration using different random seeds. DEVIL, glmGamPoi, and NEBULA were fitted using identical cell-level design matrices. Genes with low total counts were filtered prior to fitting, and end-to-end runtimes were recorded.

#### Evaluation of batch-effect control

Datasets were simulated across three orthogonal axes: study scale (small: 3,000 cells, 2 batches × 3 patients × 500 cells per patient; large: 20,000 cells, 5 batches × 4 patients × 1,000 cells per patient), batch-effect strength (low or high, jointly modulating library-size variability, gene-wise mean shifts, and dispersion across batches), and treatment effect size (weak or strong, applied to 5% of genes). In the high regime, batch effects were sufficiently strong to dominate the transcriptional structure (e.g. inducing clear batch-driven separation in low-dimensional embeddings). Ten replicates were generated per combination.

Three DEVIL specifications were compared: no batch covariates or clustering; patient-clustered sandwich estimation only; and sandwich estimation with batch fixed effects in the design matrix, to disentangle the respective contributions of explicit batch adjustment and clustered variance estimation. After filtering lowly expressed genes, differential expression was assessed at FDR 5% and compared against ground truth using MCC.

### Simulation study of experimental design structure

#### Datasets

Four real scRNA-seq datasets were used: Reed et al.^36^, Suo et al.^35^, Yazar et al.^37^, and Kumar et al.^38^. For each dataset, the six most abundant cell types were retained. Donors with fewer than 50 cells of a given cell type were excluded. Genes with mean expression below 0.1 were filtered out.

#### Simulation design

Three variables were crossed: patient number (4 or 20), differential expression probability (0.5%, 2.5%, or 5% of 1,000 genes), and treatment assignment strategy. In cell-wise settings, treatment status was assigned randomly at the individual cell level (probability 0.5), preserving within-donor cellular heterogeneity. In patient-wise settings, the statistically more challenging scenario, treatment was assigned at the donor level (probability 0.5), inducing perfect confounding between patient and condition. Expression of DE genes was artificially altered in treated cells. Five independent replicates were run per cell type, treatment strategy, DE probability, and patient number.

#### Performance metrics

Detection accuracy was quantified by Matthews correlation coefficient (MCC)^32,33^, with Benjamini–Hochberg adjusted p-values. P-value calibration was assessed via the genomic inflation factor λ (ratio of observed to expected median χ^2^ statistic under the null; values near 1 indicate well-calibrated tests). Discriminative performance was further summarized by the area under the TPR–FDR curve, computed using the iCOBRA framework^40^. DEVIL was benchmarked against limma^5^, edgeR^4^, glmGamPoi^13^, NEBULA^14^, MAST^39^, and additional competitors (Supplementary Table S5).

### Cell-type marker detection

#### Datasets

Cell-type annotation performance was evaluated on four datasets: two human PBMC datasets (Lee et al.^44^; Hao et al.^41^) and two human liver datasets (Guilliams et al.^45^; MacParland et al.^46^).

#### Data preprocessing

Each dataset was preprocessed with the following quality control steps: cells with total counts more than 5 median absolute deviations (MADs) from the median, fewer than 100 expressed genes, or mitochondrial gene content above 10% were removed. Genes with mean expression below 0.1 across all cells were filtered out. Data were normalized and dimensionality-reduced via PCA using the Seurat pipeline^42^, and unsupervised clustering was performed on the reduced manifold. Original study annotations were used to assign a cell type label to each cluster.

#### Marker detection and annotation

For each cluster, DE analysis was performed against all remaining cells using DEVIL, NEBULA, and glmGamPoi. Significantly up-regulated genes were defined by log₂ FC ≥ 1 and *p* < *p*_thresh_. The top *N* DE genes per cluster were passed to scMayoMap^43^ for cell-type annotation; for clusters with multiple candidate matches, the label with the highest confidence score was retained. Stability of the cell-type markers annotation accuracy was evaluated across N ∈ {5, 10, 25, 100, 250, 500, 1,000} and *p*_thresh_∈ {0.05, 0.01, 10⁻¹⁰, 10⁻²⁰}.

### Marker gene recovery in purified immune populations

#### Dataset

The Liu dataset^47^ contains 10 immune cell populations isolated by antibody-based positive and negative selection, uniquely barcoded, pooled, and sequenced. This design provides experimentally validated cell-type identities independent of computational clustering, offering an unambiguous ground truth for marker recovery evaluation.

#### Preprocessing and DE analysis

Raw count matrices were converted to gene-by-cell format; genes with mean count ≤ 0.05 were filtered prior to analysis. DE was performed directly using the known labels without clustering. For each cell type, a binary contrast (target population vs. all others) was fitted using DEVIL, glmGamPoi, and NEBULA with identical model specifications.

#### Performance evaluation

For each method and cell type, genes were ranked by differential expression significance and effect size. Ranked lists were compared to curated immune cell-type marker annotations from PanglaoDB^66^, with synonymous labels harmonized across datasets. ROC curves were constructed using marker membership as binary ground truth and DE scores as predictors; AUC values were computed per cell type and aggregated across populations to compare overall performance and variability between methods.

### Skeletal muscle case study

#### Data preprocessing

snRNA-seq count data from 22 individuals (age 15–99) were downloaded from the Human Muscle Ageing Cell Atlas (HLMA)^48^. The original matrix comprised 212,774 nuclei and 60,609 genes. We retained only Type I and Type II myofiber cells and applied the following quality control filters: cells with total counts exceeding the median plus 5 MADs, fewer than 100 expressed genes, or a number of expressed genes exceeding 5 MADs were removed. Genes with mean count below 0.01 across all cells were excluded. The final dataset comprised 19 patients, 126,124 nuclei, and 16,489 genes. Individuals were stratified into young (< 45 years) and old (≥ 45 years) groups for downstream analysis.

#### DE analysis

DE analysis was performed with DEVIL, NEBULA, and glmGamPoi. For each method, we fitted a linear model with main effects for age group and myofiber subtype and their interaction: expression ∼ age + cell type + age × cell type. This formulation enabled inference of (i) age effects within each subtype and (ii) cell-type-specific ageing effects via the interaction term. Size factors were computed from total cell counts. DEVIL was fitted with patient-level clustering to account for within-individual correlation; NEBULA incorporated patient-specific random effects; glmGamPoi was run without multi-sample correction. Differential expression testing was performed using contrast vectors corresponding to the main age effects within each cell type and to the interaction term. After testing, *p*-values were adjusted using the Benjamini–Hochberg procedure. Genes were considered DE at adj-*p* < 0.05 and |log₂ FC| > 1.0.

#### Functional enrichment analysis

GSEA was performed using the gseGO function from clusterProfiler^67^ on GO biological process (BP) terms (gene set size between 10 and 350). Genes were ranked by *R*(*g*) = − log_10_(*p*-value_*g*_) ⋅ sign(log_2_ *FC*). Statistical significance was assessed with fgsea (50,000 permutations) and Benjamini–Hochberg correction; GO terms with q < 0.05 were retained. GSEA was performed separately for the Type I age effect, Type II age effect, and age-by-cell-type interaction contrasts.

#### GO semantic embedding

To compare pathway-level interpretations across methods beyond discrete term lists, we embedded enriched Gene Ontology (GO) biological process (BP) terms into a two-dimensional semantic similarity space. Significantly enriched GO BP terms (p < 0.05) from each method and contrast were pooled, retaining unique GO identifiers across methods and contrasts. A pairwise semantic similarity matrix *S* was computed using GOSemSim^68^ with the Wang graph-based measure. Missing similarities were set to zero and similarities were then converted to dissimilarities (*D* = 1 − *S*) and embedded in two dimensions. Each GO term is represented as a point with size proportional to −log₁ ₀ (adj-*p*); a minimum spanning tree was overlaid within each method to aid interpretation of local structure.

#### Tissue-specificity classification

For each significantly enriched GO term, core enrichment genes were extracted and subjected to tissue-enrichment analysis using TissueEnrich^69^, based on Human Protein Atlas expression data. For each GO term, the tissue with the highest fold-change among those reaching significance (p < 0.05) was assigned; ties were broken by the number of tissue-specific genes. If no tissue met this criterion, we assigned a generic label. This yielded a GO-term–tissue pairing per method, summarized as the fraction of terms assigned to each tissue.

#### Subsampling and robustness analysis

To evaluate robustness to uneven cell representation, the full DE and GSEA pipeline was repeated on subsampled datasets equalizing cell numbers across age groups and myofiber subtypes. Subsampling was performed independently within each patient to preserve the multi-sample structure. Stability was assessed by the overlap between GO term sets from full and subsampled data.

### GPU acceleration

DEVIL’s GPU-accelerated implementation offloads expression coefficient initialization and inference, followed by overdispersion estimation, to one or more GPUs. These two steps dominate the overall computational cost of model fitting and represent the primary scalability bottlenecks.

#### Design principles

The implementation is designed to scale both runtime and the maximum tractable problem size (cells × genes). This is achieved by exploiting the conditional independence of individual ***β***_*g*_ coefficients and *θ*_*g*_ estimates, which allows inference to proceed in gene batches, accommodating GPUs with modest VRAM. The batch structure enables concurrent processing across multiple GPUs, with wall-clock time scaling approximately linearly with device count. Empirically, a batch size of 100 genes provides an optimal balance between parallelization overhead, load distribution, and memory footprint, requiring less than 10 GB of GPU memory per device.

#### GPU Expression coefficient inference

The initialization of the ***β***_*g*_ coefficients, performed before the iterative inference procedure, is designed for efficient GPU acceleration. Specifically, the coefficients are initialized by setting 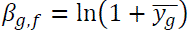 for the intercept and β_*g*,*f*_ = 0 otherwise. This approach relies solely on calculating the row averages of the input matrix, an operation that is highly parallelizable and computationally efficient on GPUs, due to their high memory bandwidth. The update equations are implemented using the cuTENSOR library^70^, which permits reformulation as three-dimensional tensor products, enabling coalesced memory access and mixed-precision computation. We use 32-bit floating-point for input and output, with 64-bit intermediates where required for numerical stability. Matrix inversions, arising at each update step to obtain ***Σ***_***β***_ estimates, are handled via batched LU factorization with partial pivoting using cuBLAS^71^, optimized for small matrices (*n* ≤ 32), with kernel launch overhead minimized as much as possible. Specifically, for the problem at hand, typical matrix sizes to be inverted are usually less than 10. The backend takes advantage of these small sizes by using specialized kernels, confirmed through profiling, which are tailored to the data type and matrix dimensions to extract maximum performance from the hardware.

#### GPU Overdispersion inference

Following convergence of ***β***_*g*_ coefficients, the overdispersion parameter *θ*_*g*_ is estimated using the method-of-moments (MoM) estimator (Methods). The MoM formulation relies entirely on Level-2 and Level-3 BLAS^72^ operations and element-wise reductions, supplemented by custom CUDA kernels for necessary element-wise computations. This, together with the ***β***_*g*_ inference, allows us to perform all computations on GPUs, reducing CPU compute time to a minimum. Full details of the GPU implementation, including pseudocode and algorithmic choices, are provided in Supplementary Information.

#### Parallelization

Finally, we implemented multi-GPU scaling by spawning one thread per device to process batches of coefficients in parallel. While larger gene batches reduce parallelization overhead, they can introduce load imbalance, as specific genes require more iterations to converge. Empirically, we found that a batch size of 100 provides an optimal balance between computational overhead, load distribution, and memory constraints.

#### High-performance computing environment

All experiments were performed on the ORFEO HPC cluster. GPU benchmarks used nodes equipped with either 8× NVIDIA A100 or 8× NVIDIA H100 accelerators. CPU-only benchmarks were run on the same H100 nodes to ensure a fair comparison. The software environment comprised CUDA 12.6, cuBLAS 12.6, and cuTENSOR 2.2.0.0. CPU benchmarks used R 4.3.3 with OpenBLAS 0.3.29 as the linear algebra backend. All software components, including DEVIL, R, and OpenBLAS, were compiled from source on the target architecture to maximize performance. Detailed hardware specifications are provided in Supplementary Table S10.

## Data Availability

No new data were generated during the study. The data that support the findings of this study are all publicly available. The accession codes for all the analyzed datasets are available in the README file available at https://zenodo.org/records/20068224 (ref.^73^)

## Code Availability

The DEVIL framework is implemented in R and is available as an open-source R/C++ package at https://github.com/caravagnalab/devil and the frozen version used for the analysis is available at https://zenodo.org/records/20068507 (ref.^74^). The code to replicate the analysis of this paper, along with data accession links, are available at https://zenodo.org/records/20068224 (ref.^73^).

Differential expression testing was carried out using edgeR v4.2.2, limma v3.60.6, glmGamPoi v1.16.0, NEBULA v1.5.6, and MAST v1.30.0. Single-cell data processing and clustering were performed with Seurat v5.4.0 and SingleCellExperiment v1.26.0. Data manipulation and visualization relied on the tidyverse v2.0.0, including ggplot2 v4.0.2, dplyr v1.2.0, tidyr v1.3.2, and purrr v1.2.1, together with patchwork v1.3.2 and ggh4x v0.3.1 for figure assembly.

## Acknowledgments

The authors acknowledge support from the Italian Association for Cancer Research (AIRC) under grant My First AIRC Grant 2020 – ID. 24913 project (PI – Giulio Caravagna), AIRC under Bridge Grant 2025 – ID.32107 (PI – Giulio Caravagna), and Investigator Grant 2021 – ID. 27631 (PI – Guido Sanguinetti). Giulio Caravagna acknowledges financial support under the National Recovery and Resilience Plan (NRRP), Mission 4, Component 2, Investment 1.1, Call for tender No. 1409 published on 14.9.2022 by the Italian Ministry of University and Research (MUR), funded by the European Union – NextGenerationEU– CUP J53D23015060001. Leonardo Egidi acknowledges financial support under Decreto Direttoriale No. 104 published on 02-02-2022 by MUR (NextGeneration EU – CUP J53D23003860006). Stefano Cozzini acknowledges financial support from the European Union – NextGenerationEU within the NRRP project PRP@CERIC” IR0000028 – Mission 4 Component 2 Investment 3.1 Action 3.1.1.

The authors acknowledge the AREA Science Park supercomputing platform ORFEO made available for conducting the research reported in this paper, as well as the technical support of the Laboratory of Data Engineering staff.

## Author Contributions Statement

GiS, NT, SM developed the framework with the supervision of GuS, LE and GC. GiS and NT implemented the model, with the supervision of SC. GiS implemented synthetic data and tested the model with EI. GiS, NT and KD gathered and analyzed the data for the case studies. GiS, NT, SC, LE and GC drafted the manuscript, which all authors approved. SC, LE and GC conceptualized and supervised this work.

