## Supplementary Information for "Scalable, fast and accurate differential gene expression testing from millions of cells of multiple patients"

### 1 Supplementary Figures

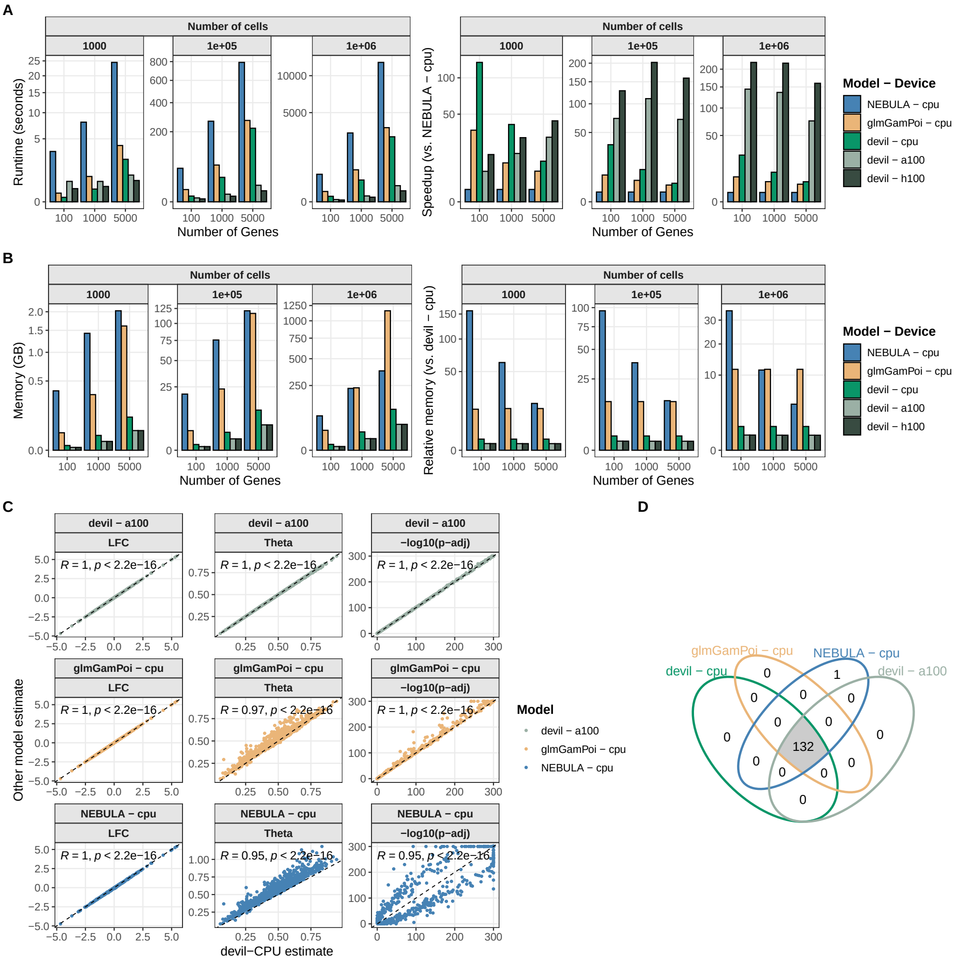


**Supplementary Figure 1** Performance comparison for the dataset presented by Chiou et al.[1]. **A** Absolute runtime comparison and speedup of devil (CPU and GPU implementations), glmGamPoi and NEBULA for the differential expression analysis of scRNA-seq data. **B** Memory consumption comparison of devil, glmGamPoi, and NEBULA for the differential expression analysis of scRNA-seq data. For every dataset size. **C** Correlation between log-fold changes (LFC, overdispersion (Theta) and adjusted p-values obtained with the different tools/versions. Points represent individual genes. Correlation was assessed using Pearson's correlation coefficient (two-sided test), with R and corresponding p values shown in each panel. **D** Number of differentially expressed genes retrieved by the different tools.

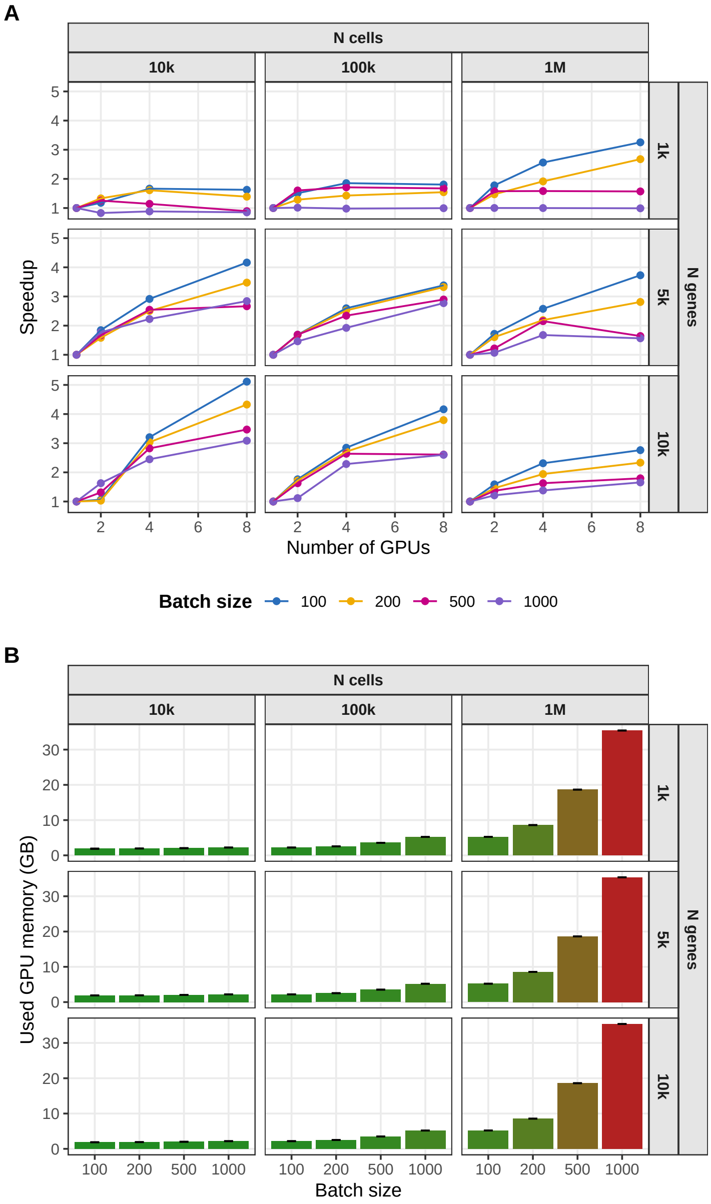


**Supplementary Figure 2 Strong scaling and GPU memory usage of devil's GPU implementation. A.** Strong-scaling benchmark showing runtime speedup as a function of the number of GPUs (1–8) for datasets spanning 10k, 100k, and 1M cells and 1k, 5k, and 10k genes, evaluated at different batch sizes. The smaller the batch size, the greater the work distribution granularity. **B.** Corresponding GPU memory usage per device as a function of batch size for the same datasets. A batch size of 100 consistently achieves near-optimal scaling while keeping memory usage below 6 GB per GPU, even for the largest datasets, enabling efficient execution even on a consumer-grade accelerator.

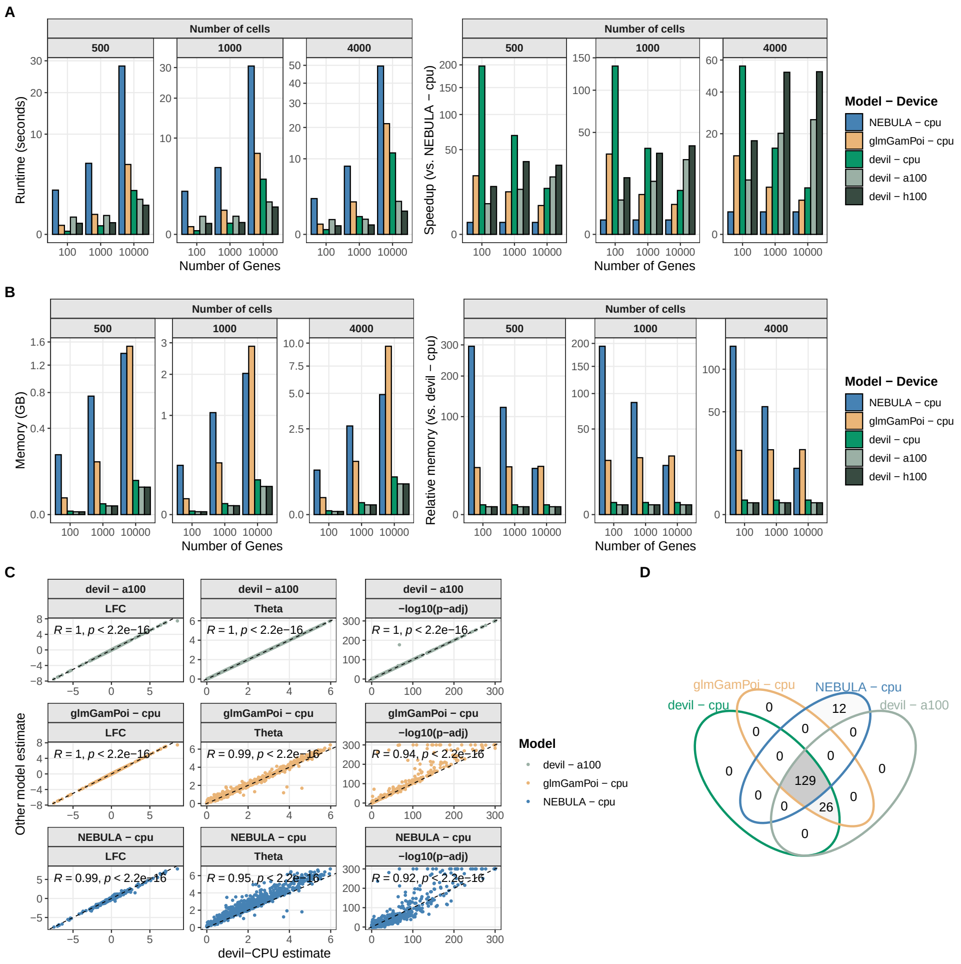


**Supplementary Figure 3** Performance comparison for the dataset presented by Baron et al.[2]. **A** Absolute runtime comparison and speedup of devil (CPU and GPU implementations), glmGamPoi and NEBULA for the differential expression analysis of scRNA-seq data. **B** Memory consumption comparison of devil, glmGamPoi, and NEBULA for the differential expression analysis of scRNA-seq data. For every dataset size. **C** Correlation between log-fold changes (LFC, overdispersion (Theta) and adjusted p-values obtained with the different tools/versions. Points represent individual genes. Correlation was assessed using Pearson's correlation coefficient (two-sided test), with R and corresponding p values shown in each panel. **D** Number of differentially expressed genes retrieved by the different tools.

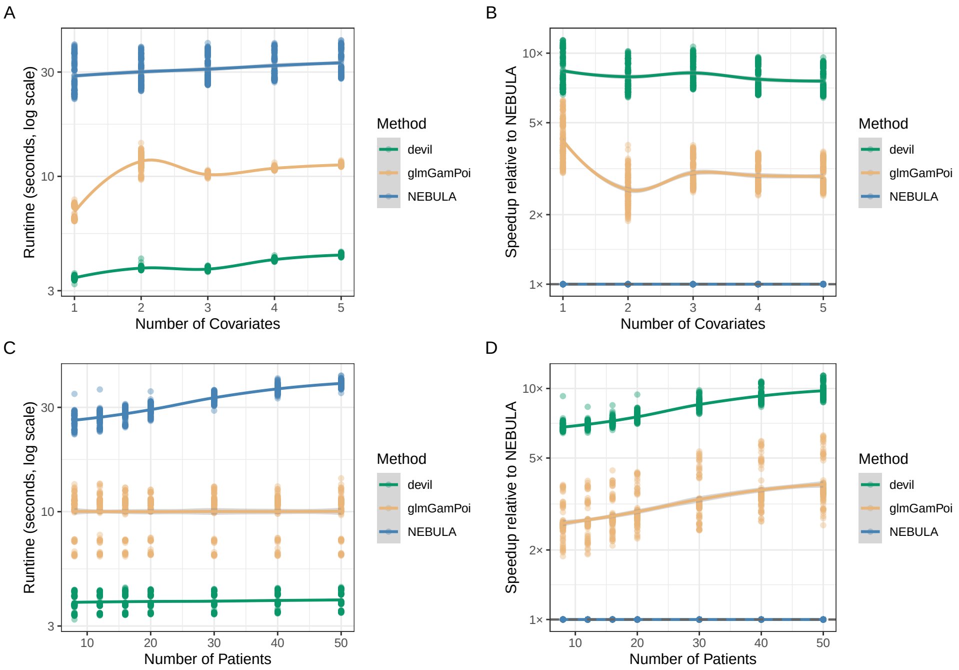


**Supplementary Figure 4 Scaling across patients and features. A.** Runtime (log scale) of devil, glmGamPoi, and NEBULA as a function of the number of covariates (1–5), with the total number of cells held constant. **A.** Corresponding speedup relative to NEBULA for each method across increasing feature dimensionality. **C.** Runtime (log scale) as a function of the number of patients (8–50), again at fixed total cell count. **D.** Corresponding speedup relative to NEBULA. Increasing the number of patients has a pronounced impact on NEBULA's runtime, while this is less evident for both devil.

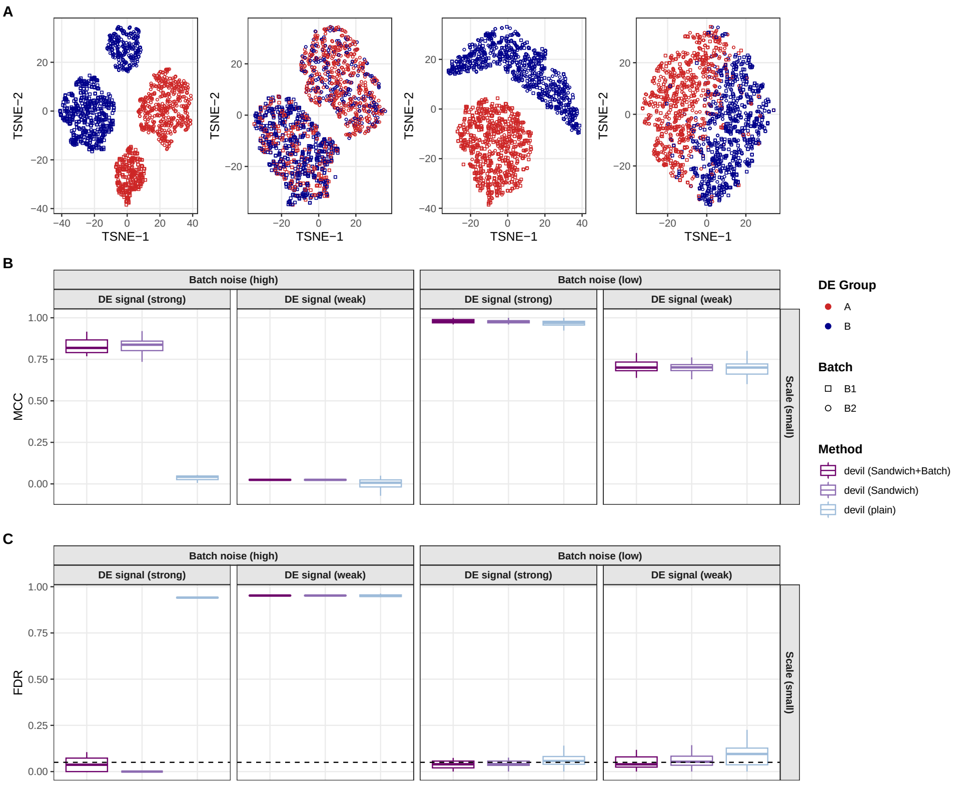


**Supplementary Figure 5 Batch-effect control benchmark at small study scale. A.** Two-dimensional t-SNE embeddings of simulated datasets under a small-scale design (2 batches, 3 patients per batch, 500 cells per patient), illustrating combinations of high vs low batch noise and strong vs weak differential expression (DE) signal. Points are coloured by treatment group (A/B) and shaped by batch, highlighting scenarios in which batch effects dominate the transcriptional structure. **B.** Matthews correlation coefficient (MCC) for treatment differential expression detection across simulation regimes, comparing three devil analysis strategies: plain (no batch adjustment, no clustering), sandwich-robust inference with patient clustering, and sandwich-robust inference with explicit batch covariates. **C** Corresponding false discovery rate (FDR) at a nominal 5% threshold. Each box-plot summarizes 10 independent simulation replicates. Boxes show the IQR with median indicated; whiskers extend to 1.5× IQR.

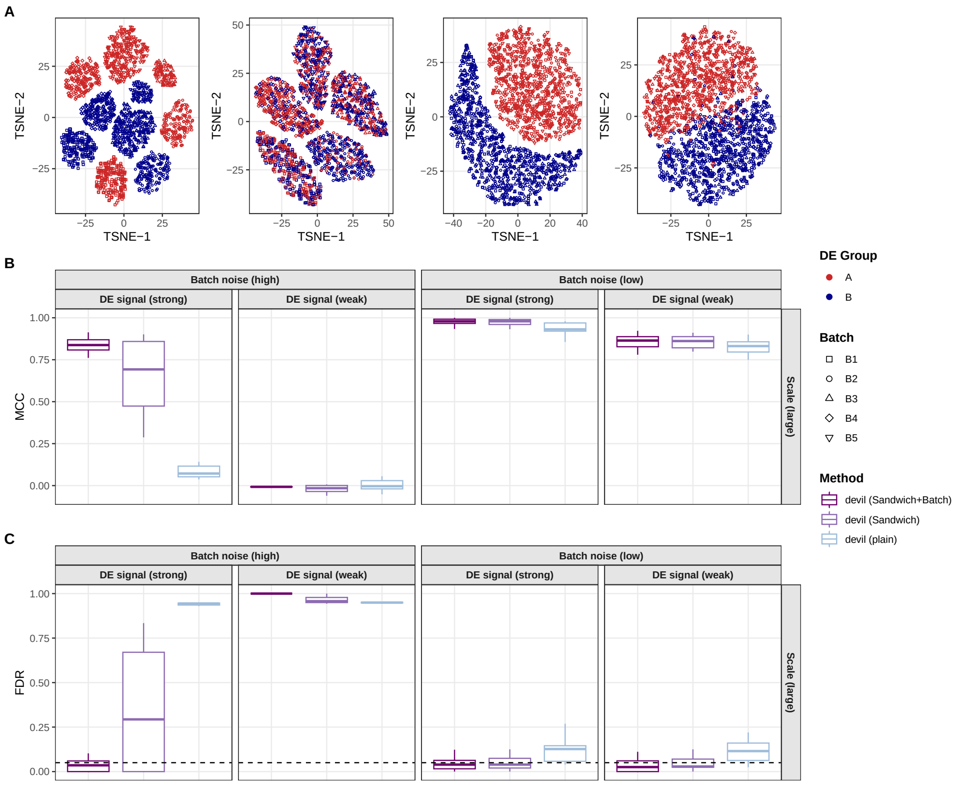


**Supplementary Figure 6  Batch-effect control benchmark at large study scale. A.** Two-dimensional t-SNE embeddings of simulated datasets under a small-scale design (5 batches, 4 patients per batch, 1,000 cells per patient), illustrating combinations of high vs low batch noise and strong vs weak differential expression (DE) signal. Points are coloured by treatment group (A/B) and shaped by batch, highlighting scenarios in which batch effects dominate the transcriptional structure. **B.** Matthews correlation coefficient (MCC) for treatment differential expression detection across simulation regimes, comparing three devil analysis strategies: plain (no batch adjustment, no clustering), sandwich-robust inference with patient clustering, and sandwich-robust inference with explicit batch covariates. **C:** Corresponding false discovery rate (FDR) at a nominal 5% threshold. Each box-plot summarizes 10 independent simulation replicates. Boxes show the IQR with median indicated; whiskers extend to 1.5× IQR.

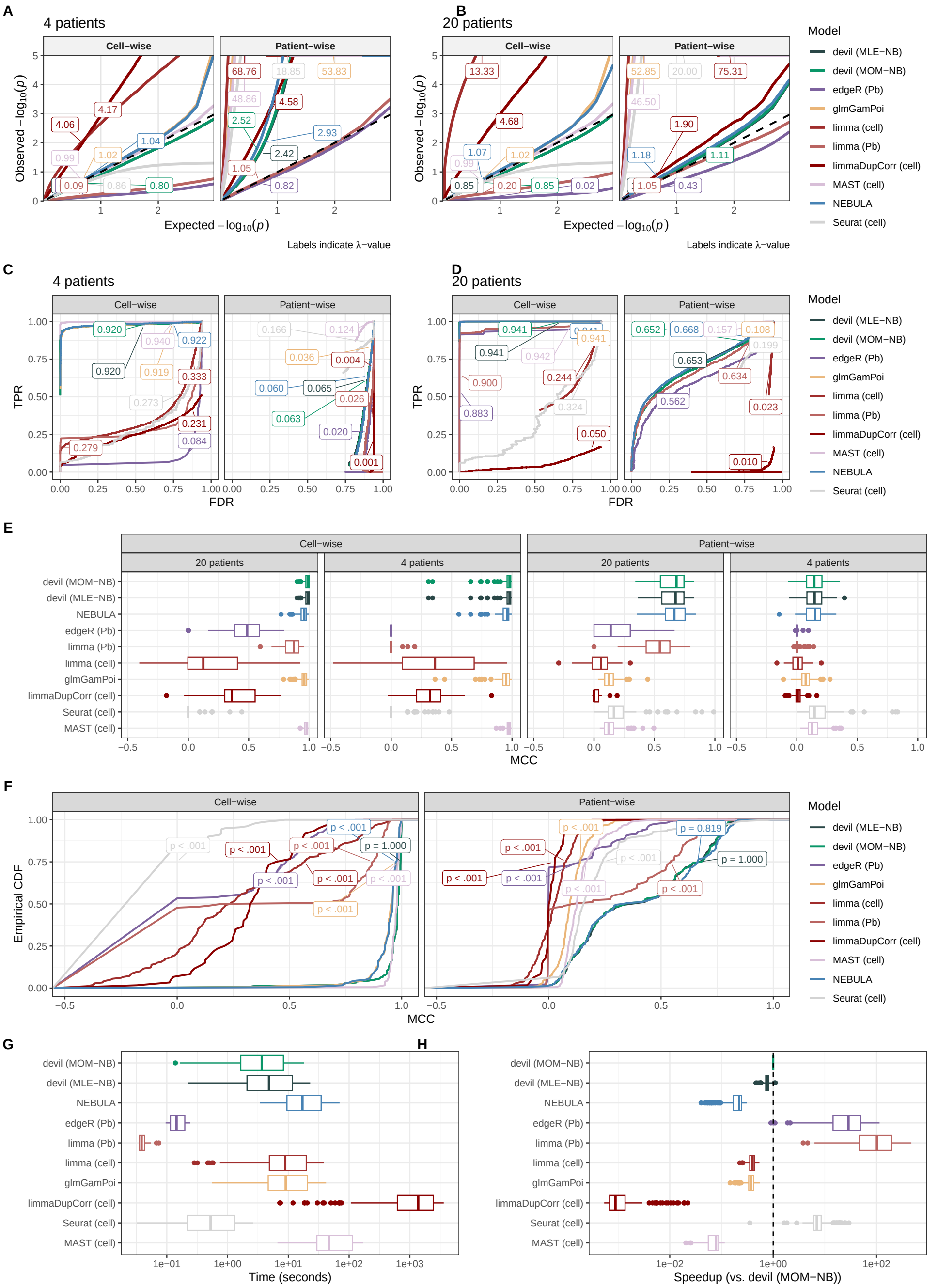


**Supplementary Figure 7** Simulation results over dataset by Reed et al.[3] Simulated datasets contain on average $N = 8,998$ cells (4-patients) and $N = 37,417$ cells (20-patients). **A-B.** QQ-plot of expected vs. observed -$\log_{10}$ p-values for non-differentially expressed genes across all tools in both experimental settings and consider 4 (A) or 20 (B) patients, with the inflation factor $\lambda$ reported for each tool. **C-D.** TPR/FDR curve computed by the iCOBRA package showing the power of a given method for a nominal FDR threshold, with AUC values reported. Both experimental settings are reported, considering 4 (C) or 20 (D) patients. **E.** Performance evaluation using Matthews Correlation Coefficient (MCC) across varying numbers of patients (4 and 20) for *cell-wise*(C) and *patient-wise*(D) settings. **F.** Empirical cumulative distribution function (ECDF) of MCC values in the cell-wise and *patient-wise* setting. Statistical comparison using Kolmogorov-Smirnov tests (p-values shown) confirms devil's performance parity with leading methods. **G,H.** Computational efficiency comparison showing total runtime (G) and time fold changes (H) relative to CPU-based devil. Boxes show the IQR with median indicated; whiskers extend to 1.5× IQR; points denote outliers.

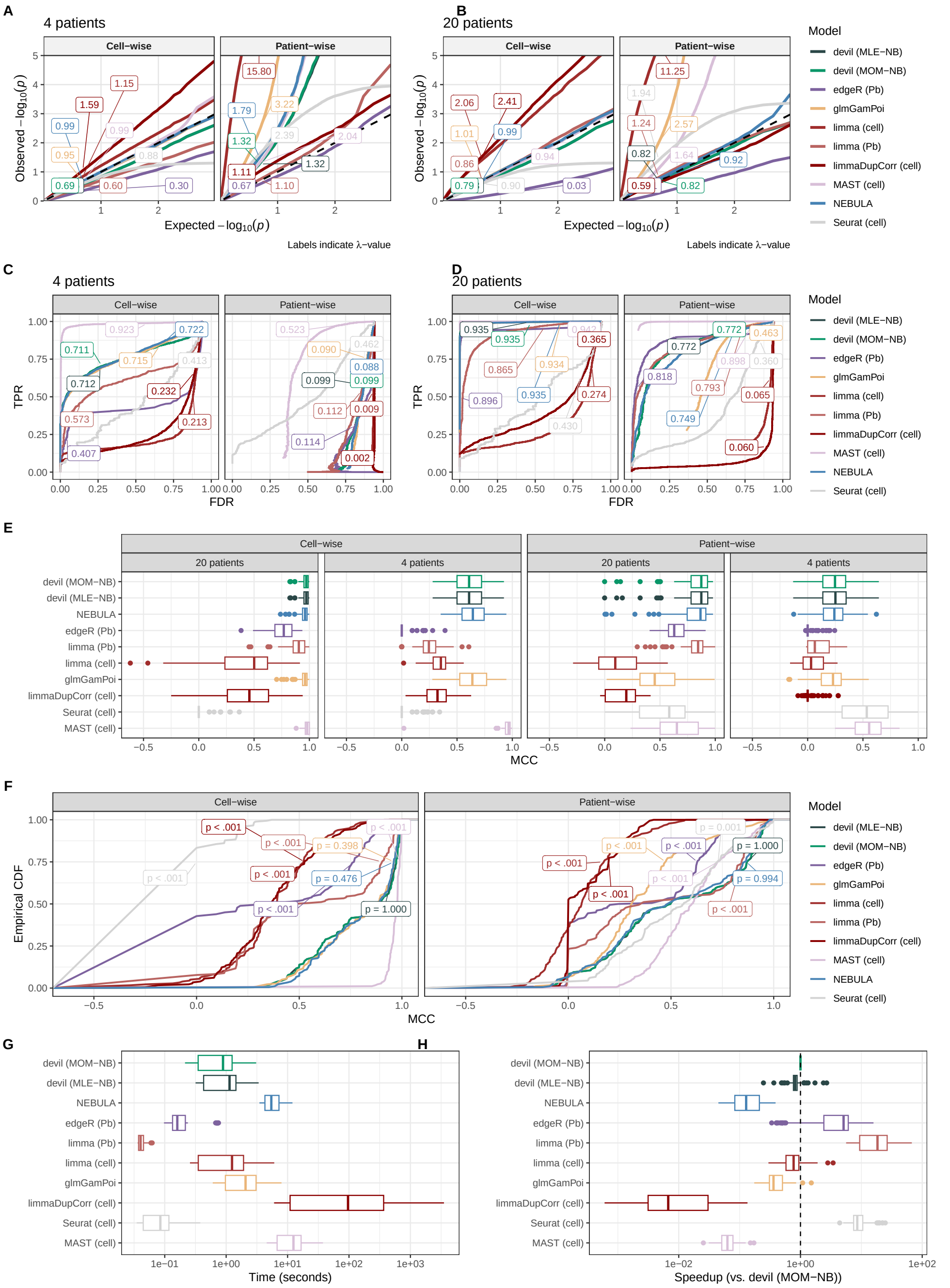


**Supplementary Figure 8** Simulation results over dataset by Yazar et al.[4]. Simulated datasets contain on average $N = 738$ cells (4-patients) and $N = 3,523$ cells (20-patients). **A-B.** QQ-plot of expected vs. observed -$\log_{10}$ p-values for non-differentially expressed genes across all tools in both experimental settings and consider 4 (A) or 20 (B) patients, with the inflation factor $\lambda$ reported for each tool. **C-D.** TPR/FDR curve computed by the iCOBRA package showing the power of a given method for a nominal FDR threshold, with AUC values reported. Both experimental settings are reported, considering 4 (C) or 20 (D) patients. **E.** Performance evaluation using Matthews Correlation Coefficient (MCC) across varying numbers of patients (4 and 20) for *cell-wise*(C) and *patient-wise*(D) settings. **F.** Empirical cumulative distribution function (ECDF) of MCC values in the cell-wise and *patient-wise* setting. Statistical comparison using Kolmogorov-Smirnov tests (p-values shown) confirms devil's performance parity with leading methods. **G,H.** Computational efficiency comparison showing total runtime (G) and time fold changes (H) relative to CPU-based devil. Boxes show the IQR with median indicated; whiskers extend to 1.5× IQR; points denote outliers.

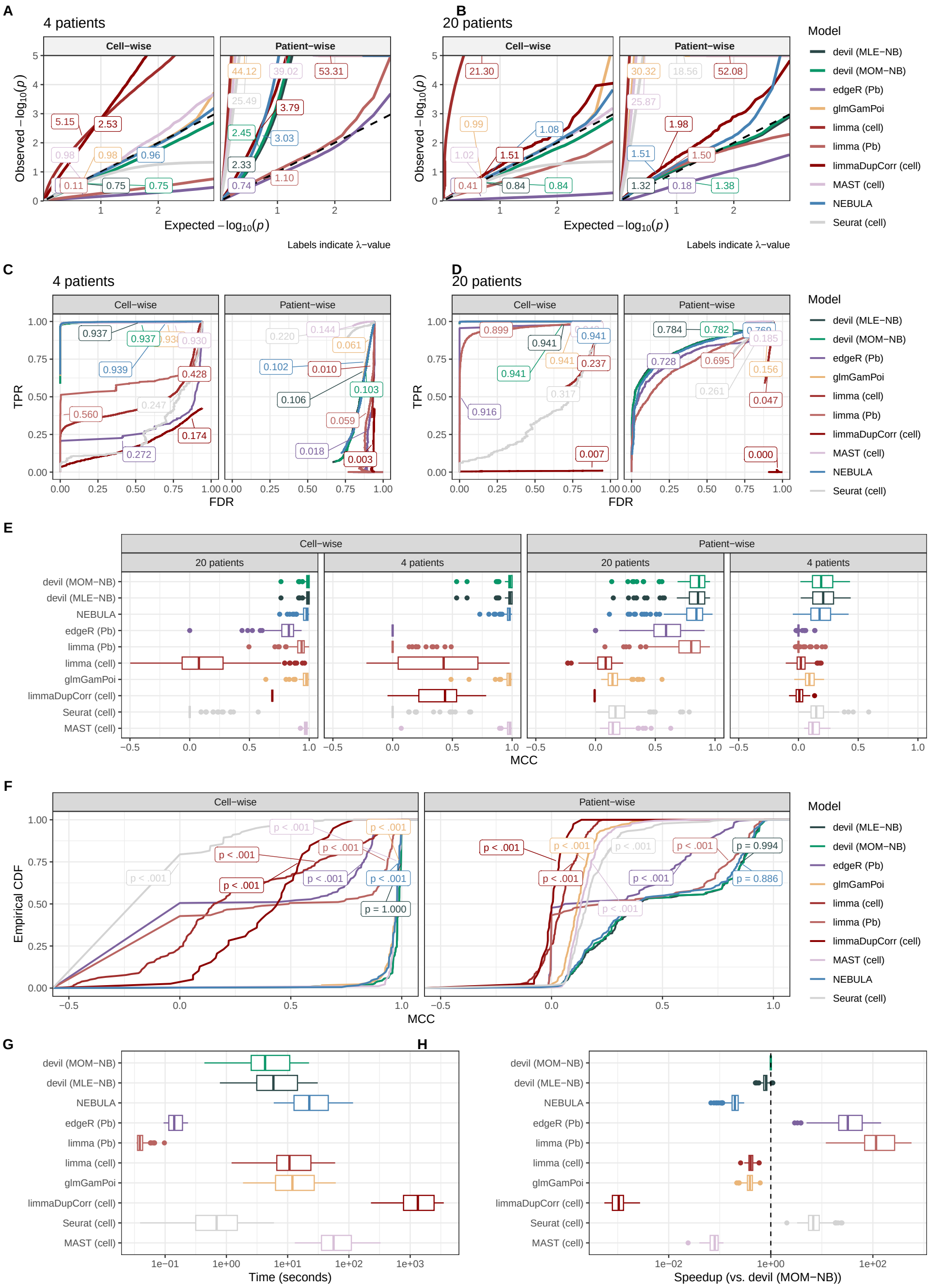


**Supplementary Figure 9** Simulation results over dataset by Suo et al.[5]. Simulated datasets contain on average $N = 9,679$ cells (4-patients) and $N = 45,294$ cells (20-patients). **A-B.** QQ-plot of expected vs. observed -$\log_{10}$ p-values for non-differentially expressed genes across all tools in both experimental settings and consider 4 (A) or 20 (B) patients, with the inflation factor $\lambda$ reported for each tool. **C-D.** TPR/FDR curve computed by the iCOBRA package showing the power of a given method for a nominal FDR threshold, with AUC values reported. Both experimental settings are reported, considering 4 (C) or 20 (D) patients. **E.** Performance evaluation using Matthews Correlation Coefficient (MCC) across varying numbers of patients (4 and 20) for *cell-wise*(C) and *patient-wise*(D) settings. **F.** Empirical cumulative distribution function (ECDF) of MCC values in the cell-wise and *patient-wise* setting. Statistical comparison using Kolmogorov-Smirnov tests (p-values shown) confirms devil's performance parity with leading methods. **G,H.** Computational efficiency comparison showing total runtime (G) and time fold changes (H) relative to CPU-based devil. Boxes show the IQR with median indicated; whiskers extend to 1.5× IQR; points denote outliers.

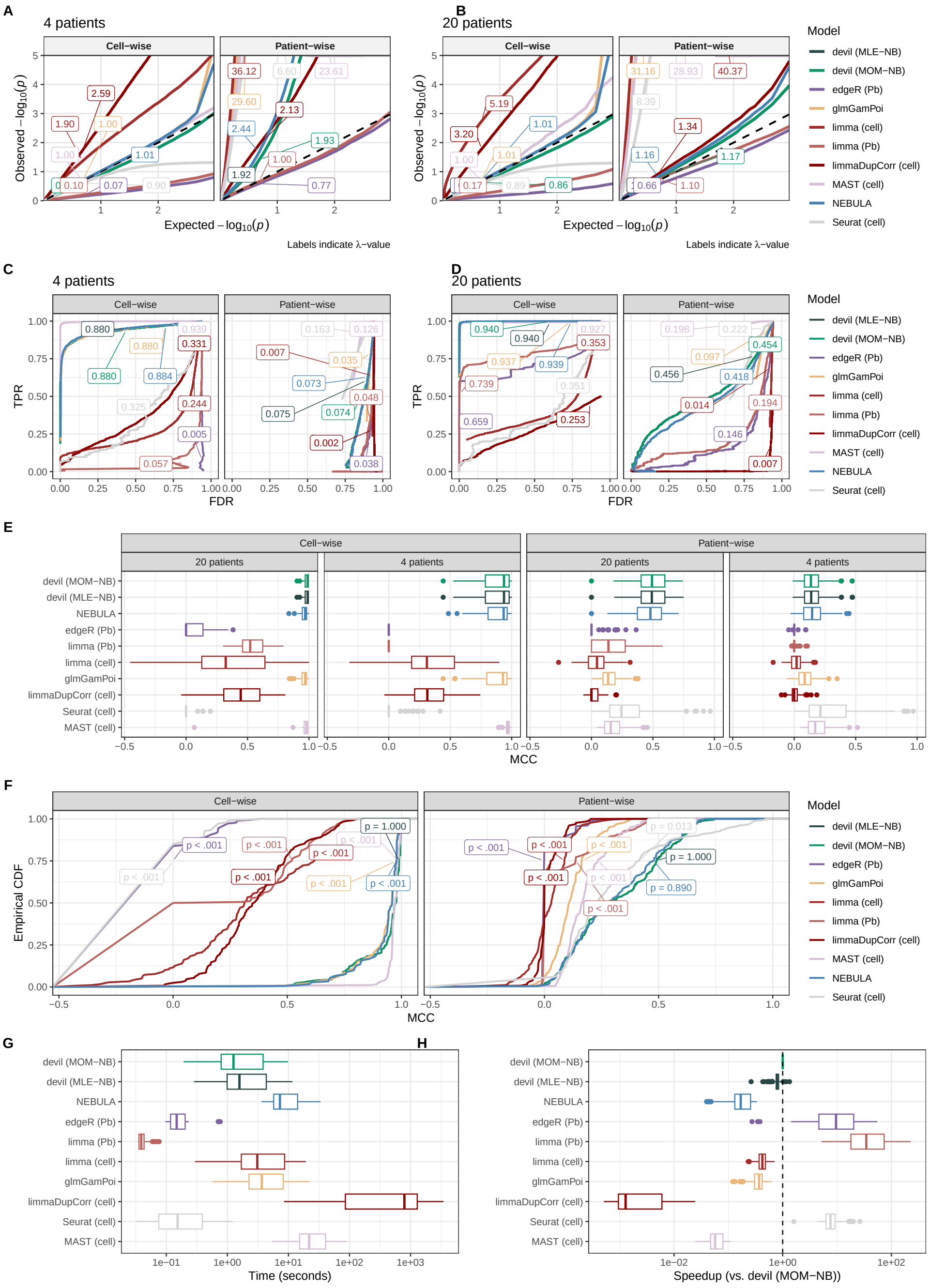


**Supplementary Figure 10** Simulation results over dataset by Kumar et al.[6]. Simulated datasets contain on average $N = 3,357$ cells (4-patients) and $N = 13,827$ cells (20-patients). **A-B.** QQ-plot of expected vs. observed -$\log_{10}$ p-values for non-differentially expressed genes across all tools in both experimental settings and consider 4 (A) or 20 (B) patients, with the inflation factor $\lambda$ reported for each tool. **C-D.** TPR/FDR curve computed by the iCOBRA package showing the power of a given method for a nominal FDR threshold, with AUC values reported. Both experimental settings are reported, considering 4 (C) or 20 (D) patients. **E.** Performance evaluation using Matthews Correlation Coefficient (MCC) across varying numbers of patients (4 and 20) for *cell-wise*(C) and *patient-wise*(D) settings. **F.** Empirical cumulative distribution function (ECDF) of MCC values in the cell-wise and *patient-wise* setting. Statistical comparison using Kolmogorov-Smirnov tests (p-values shown) confirms devil's performance parity with leading methods. **G,H.** Computational efficiency comparison showing total runtime (G) and time fold changes (H) relative to CPU-based devil. Boxes show the IQR with median indicated; whiskers extend to 1.5× IQR; points denote outliers.

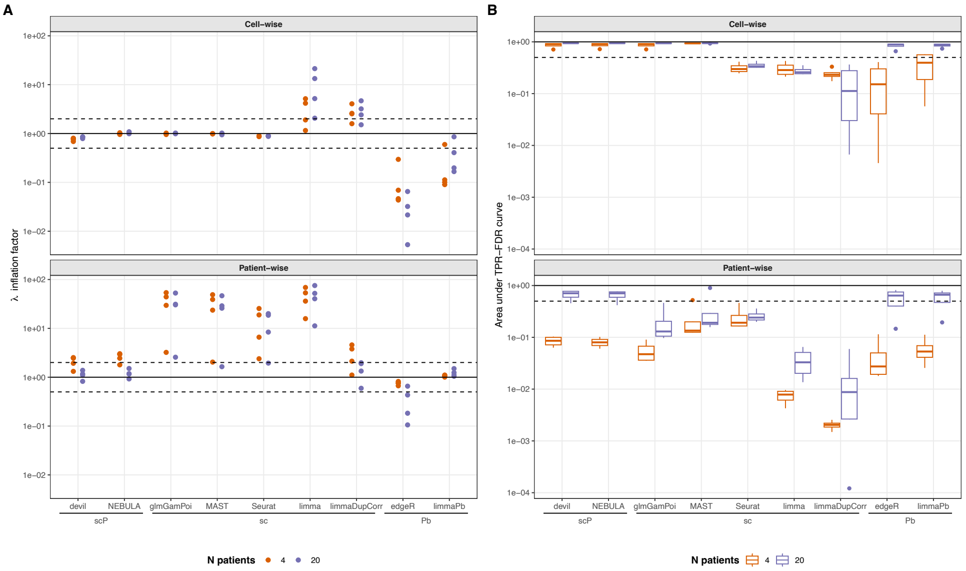


**Supplementary Figure 11** Summary of calibration and power across all benchmarked methods, datasets, and experimental settings. Each point represents one dataset (Reed et al.[3], Suo et al.[5], Yazar et al.[4], Kumar et al.[6]). Methods are grouped by class: patient-aware single-cell models (scP), standard single-cell models (sc), and pseudobulk approaches (Pb). **A.** Inflation factor $\lambda$ computed from QQ-plots of p-values under the null hypothesis, for each method across cell-wise and patient-wise experimental designs with 4 and 20 patients. Values close to 1 indicate well-calibrated tests; values substantially above 1 indicate inflated type-I error rates. **B.** Area under the TPR–FDR curve summarising detection power, under the same experimental conditions as in (A). Boxes show the IQR with median indicated; whiskers extend to 1.5× IQR.

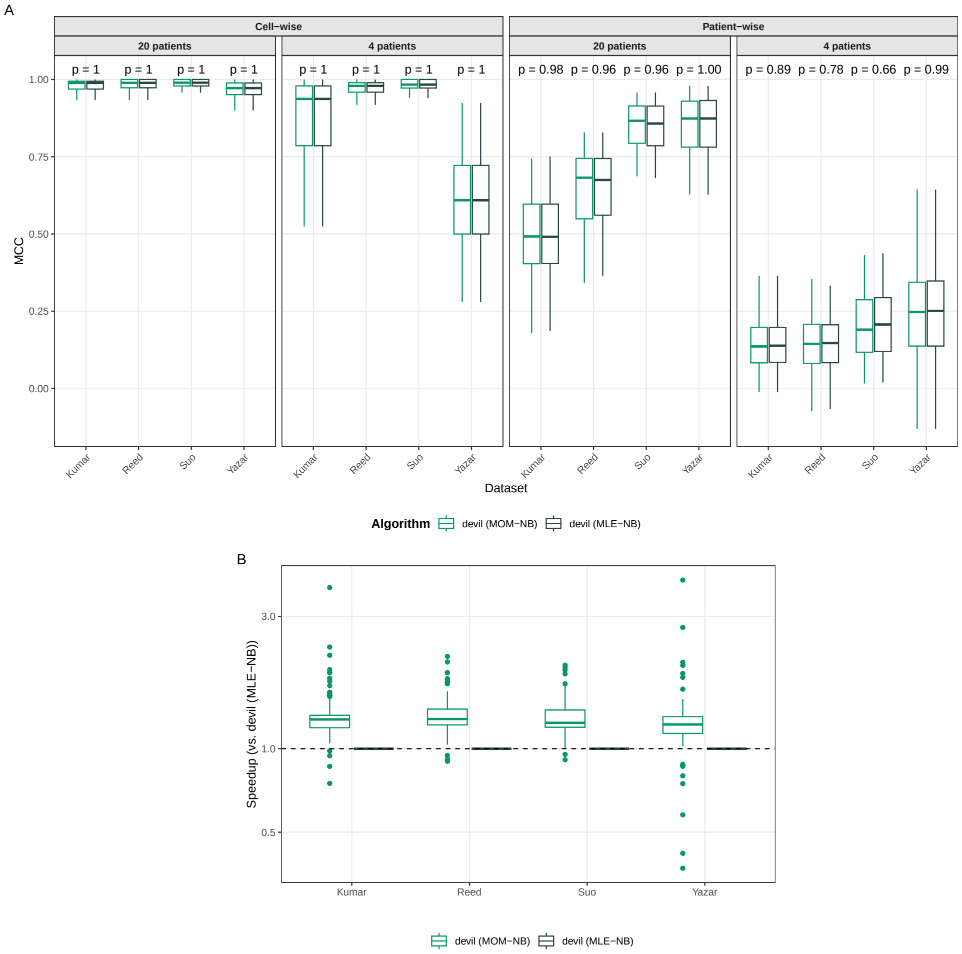


**Supplementary Figure 12** Comparison between MLE and MOM overdispersion estimators in devil. **A.** Statistical performance assessed using the Matthews Correlation Coefficient (MCC) across multiple datasets, comparing scenarios with 4 and 20 patients in both cell-wise and patient-wise settings (each combination contains n=90 simulations). Statistical comparisons were performed using two-sided Wilcoxon rank-sum tests (unpaired). P values are shown; no multiple testing correction was applied. **B.** Computational efficiency, reported as relative speedup of the MOM estimator compared to MLE. Boxes show the IQR with median indicated; whiskers extend to 1.5× IQR.

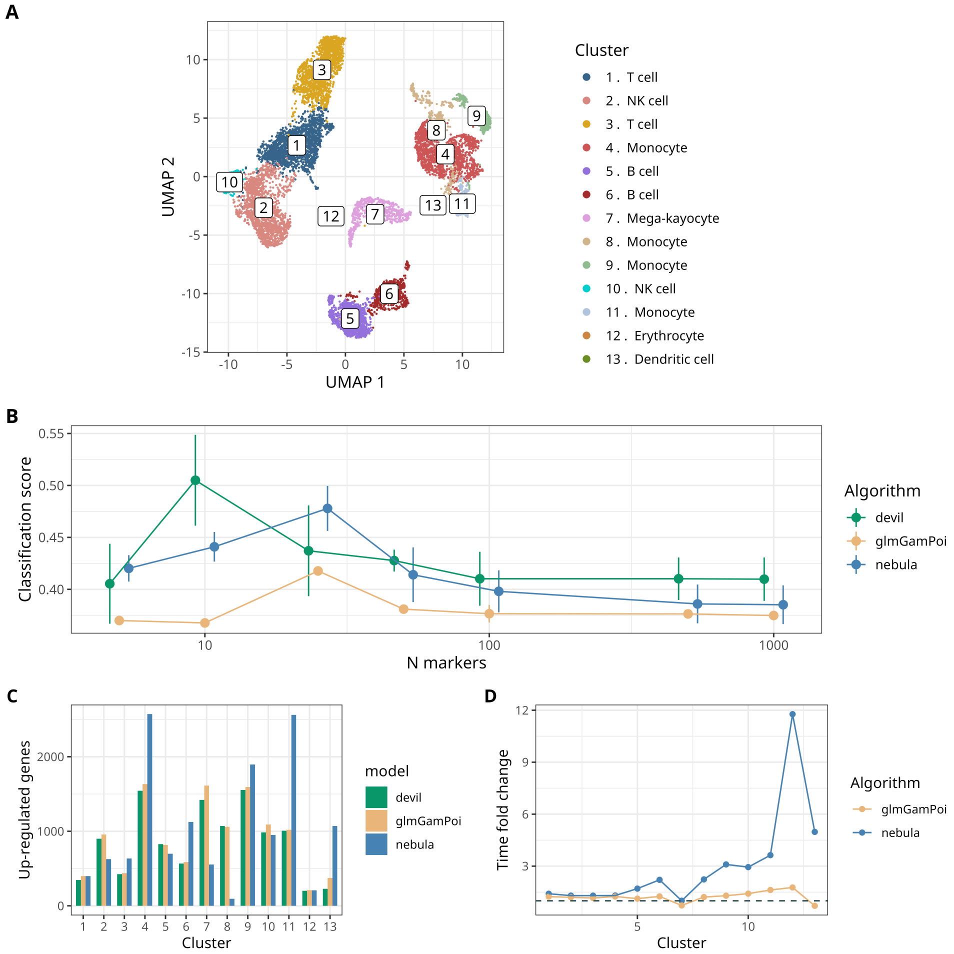


**Supplementary Figure 13** Dataset by Lee et al. [7] **A** UMAP plot of the input dataset, reporting both the clusters retrieved by Seurat and the pre-existing cell-types annotation **B.** Classification score for the tools considered with respect to the number of markers selected (x-axis) and the value of $p_{thresh}$ (error-bars). Points show mean accuracy, and error bars denote ±1 s.d. across n=4 p-value thresholds to reflect parameter sensitivity of the analysis. **C.** Numbers of up-regulated genes called by the tools in the different clusters. **D.** Timing comparison between the tools.

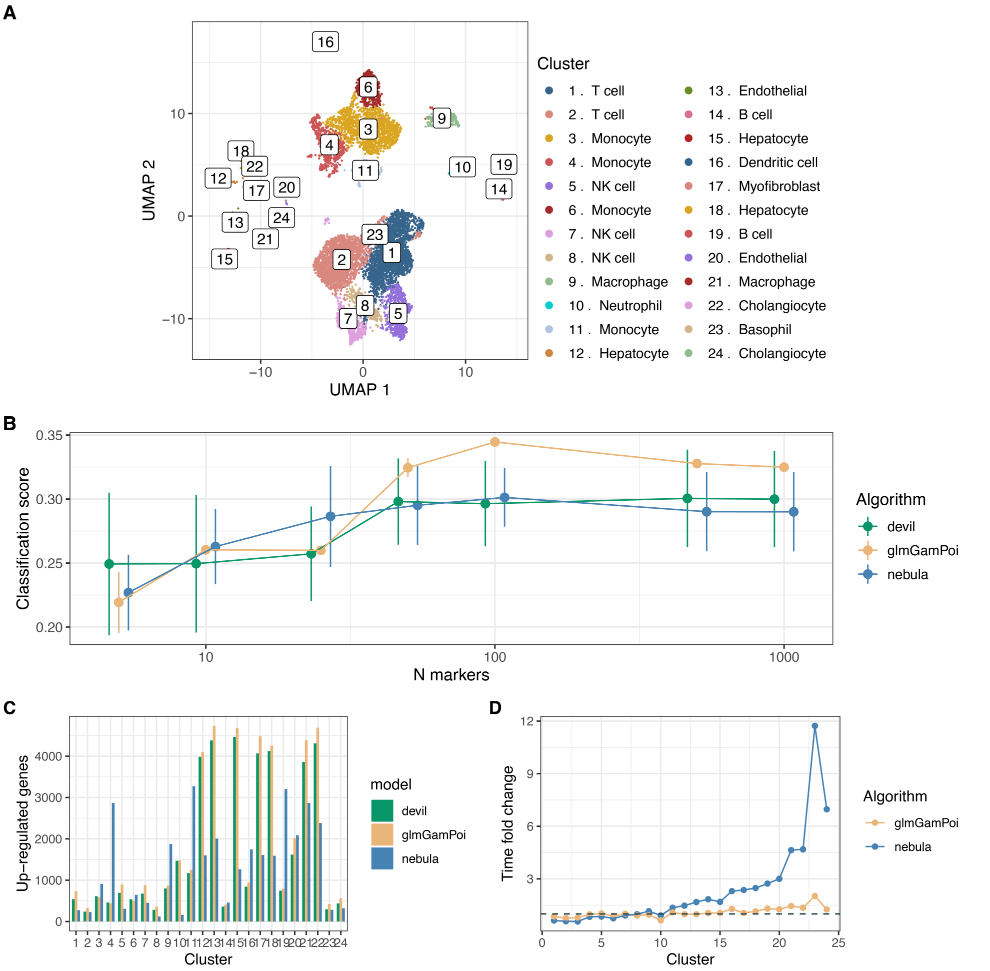


**Supplementary Figure 14** Dataset by Guilliams et al. [8] **A** UMAP plot of the input dataset, reporting both the clusters retrieved by Seurat and the pre-existing cell-types annotation **B.** Classification score for the tools considered with respect to the number of markers selected (x-axis) and the value of $p_{thresh}$ (error-bars). Points show mean accuracy, and error bars denote ±1 s.d. across n=4 p-value thresholds to reflect parameter sensitivity of the analysis. **C.** Numbers of up-regulated genes called by the tools in the different clusters. **D.** Timing comparison between the tools.

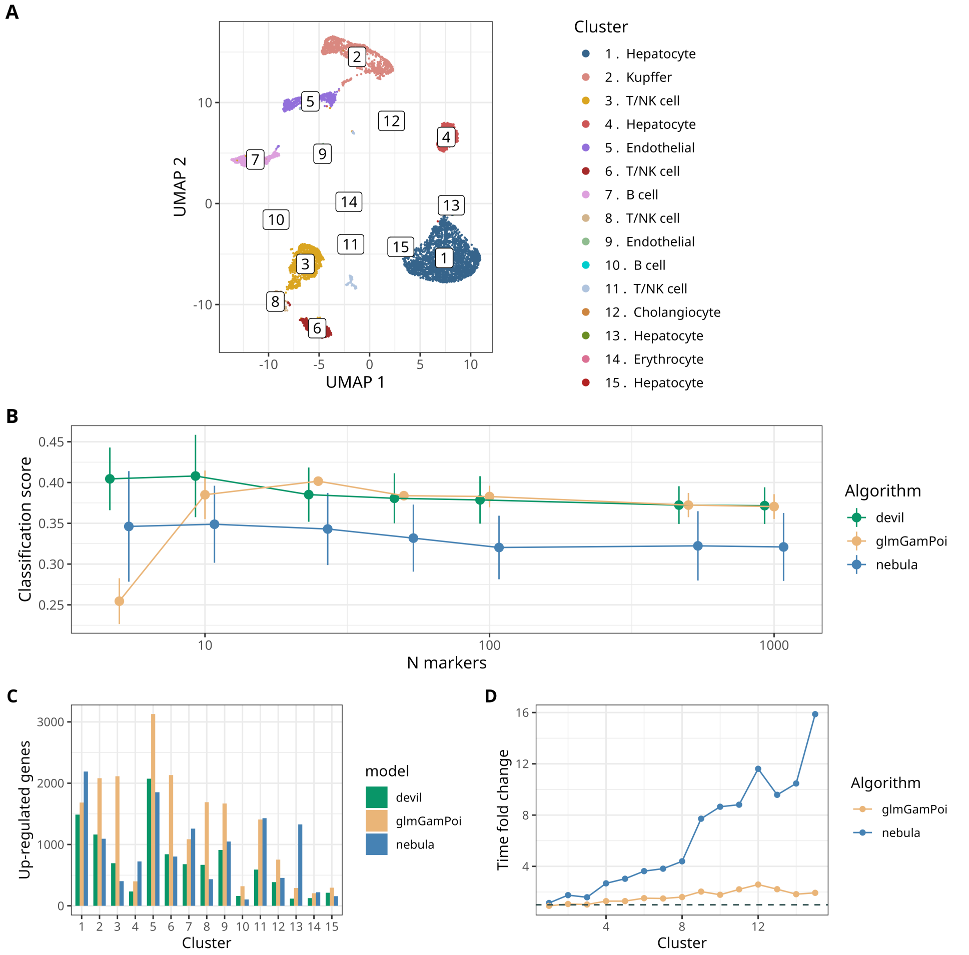


**Supplementary Figure 15** Dataset by MacParland et al. [9] **A** UMAP plot of the input dataset, reporting both the clusters retrieved by Seurat and the pre-existing cell-types annotation **B.** Classification score for the tools considered with respect to the number of markers selected (x-axis) and the value of $p_{thresh}$ (error-bars). Points show mean accuracy, and error bars denote $\pm 1$ s.d. across n=4 p-value thresholds to reflect parameter sensitivity of the analysis. **C.** Numbers of up-regulated genes called by the tools in the different clusters. **D.** Timing comparison between the tools.

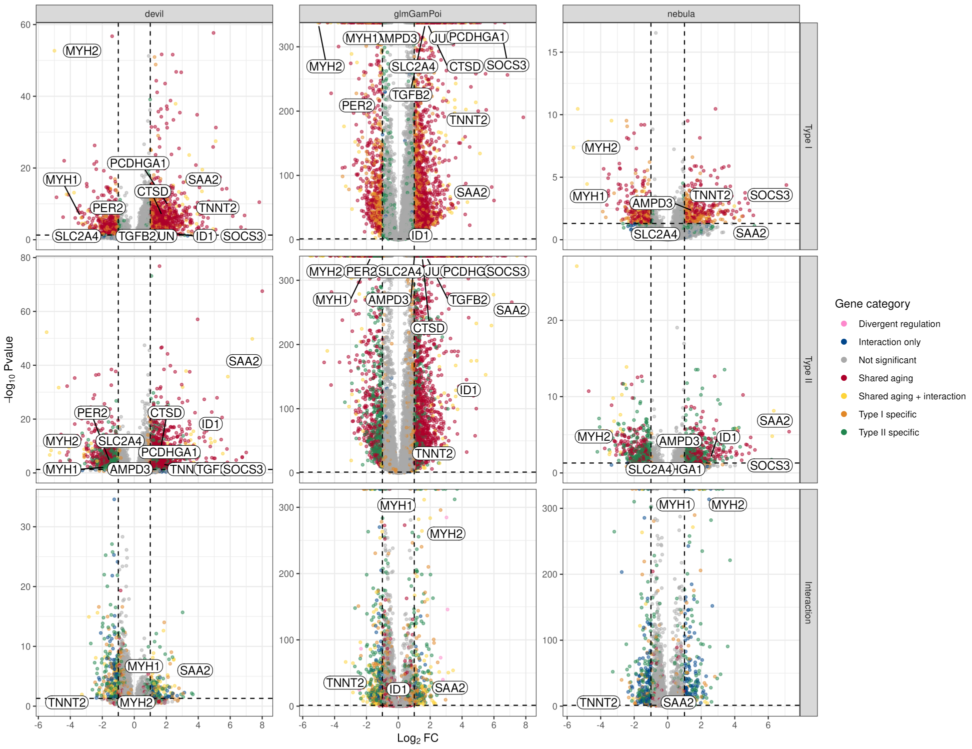


**Supplementary Figure 16 Volcano plots of ageing-associated differential expression across methods.** Volcano plots showing differential expression results from devil, glmGamPoi, and NEBULA for Type I myonuclei (top), Type II myonuclei (middle), and the age-by-myofiber subtype interaction (bottom). Each point represents a gene. Vertical dashed lines indicate fold-change thresholds and the horizontal dashed line denotes the significance cutoff. P-values were Benjamini–Hochberg adjusted. Genes are coloured by regulatory category, and selected marker genes are annotated.

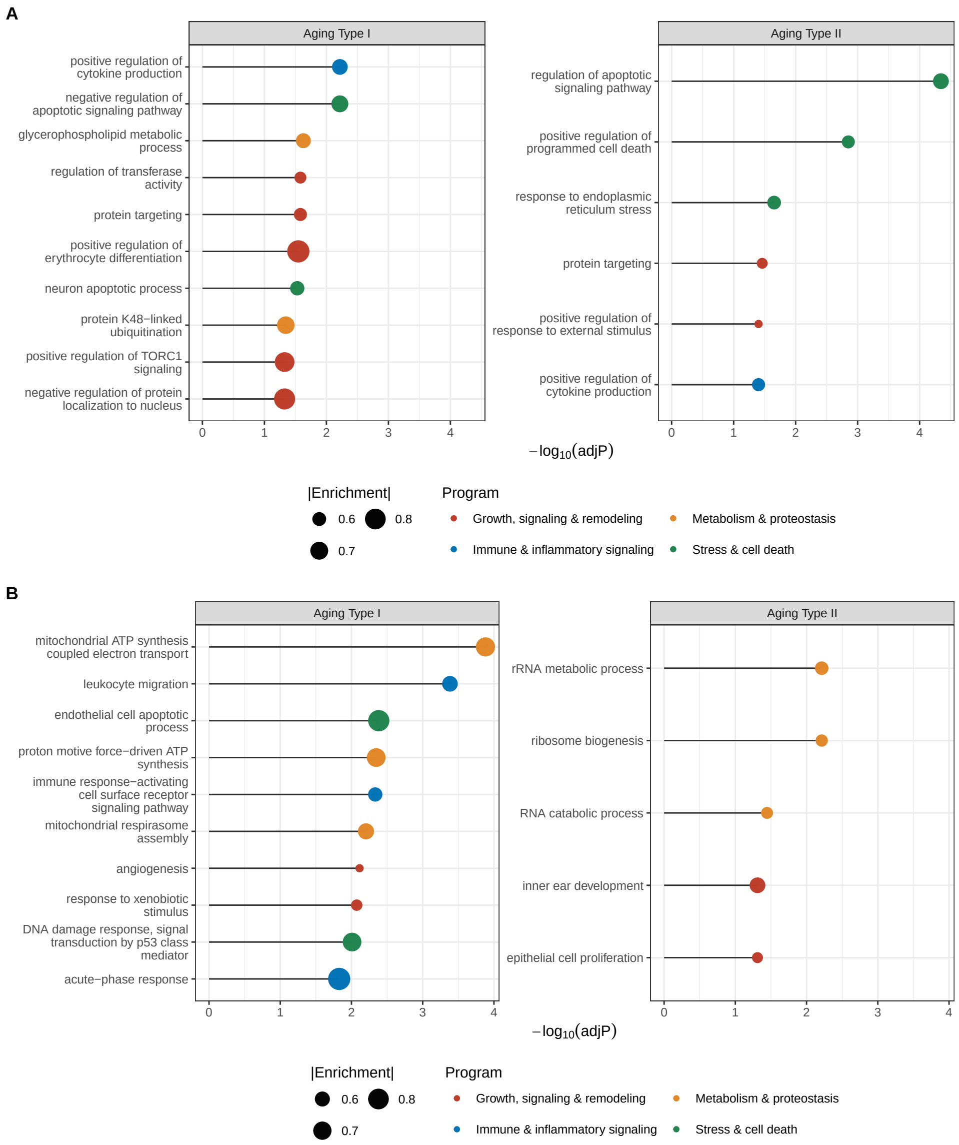


**Supplementary Figure 17 Top GSEA pathways identified by glmGamPoi and NEBULA. A,B.** Dot plots showing the top 10 significantly enriched Gene Ontology biological process terms identified by glmGamPoi (A) and NEBULA (B) for ageing contrasts in Type I and Type II myonuclei. Terms are ranked by adjusted p-value. Adjusted P values were obtained from Gene Set Enrichment Analysis (GSEA; permutation-based test with 50,000 permutations) and corrected for multiple comparisons using the Benjamini–Hochberg procedure. Dot size reflects enrichment magnitude, and colours denote broad biological program categories.

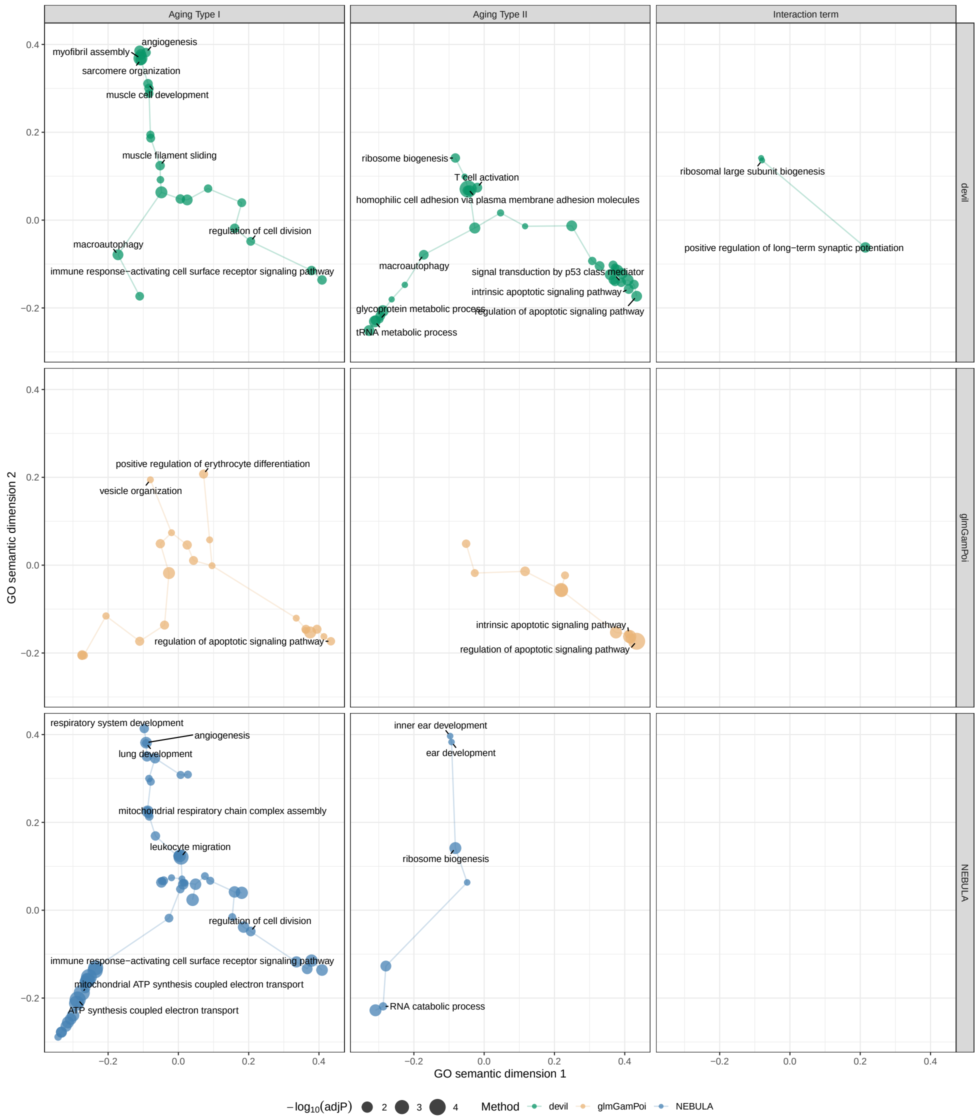


**Supplementary Figure 18 GO semantic similarity maps across methods and contrasts.** Grid of two-dimensional GO semantic embeddings of enriched biological process terms, with methods shown along the x-axis (devil, glmGamPoi, NEBULA) and contrasts along the y-axis (Type I ageing, Type II ageing, and age-by-myofiber subtype interaction). Each point represents a GO term positioned according to pairwise semantic similarity, with point size and colour indicating statistical significance. Adjusted P values were obtained from Gene Set Enrichment Analysis (GSEA; permutation-based test with 50,000 permutations) and corrected for multiple comparisons using the Benjamini–Hochberg (BH) procedure. Labels denote selected representative terms.

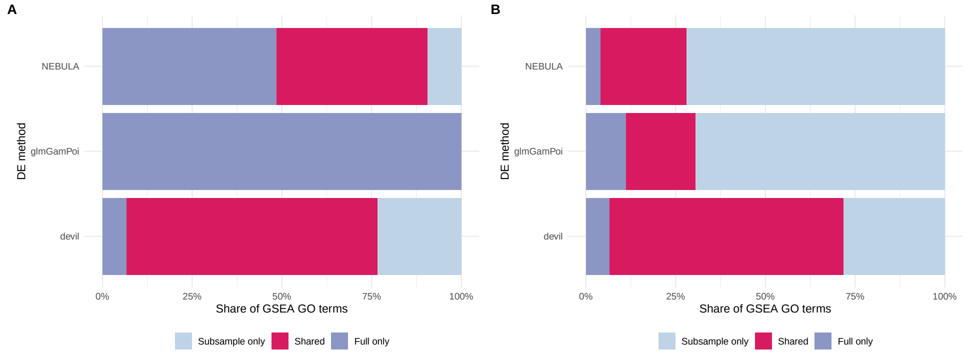


**Supplementary Figure 19 Overlap of GSEA terms between full and subsampled datasets. A,B.** Stacked bar plots showing the proportion of enriched GO biological process terms identified exclusively in the full dataset, exclusively in subsampled datasets, or shared between the two, for devil, glmGamPoi, and NEBULA. Results are shown separately for Type I (A) and Type II (B) myonuclei. Bars represent the relative share of GO terms in each category, illustrating the robustness of pathway detection to cell-number imbalance.

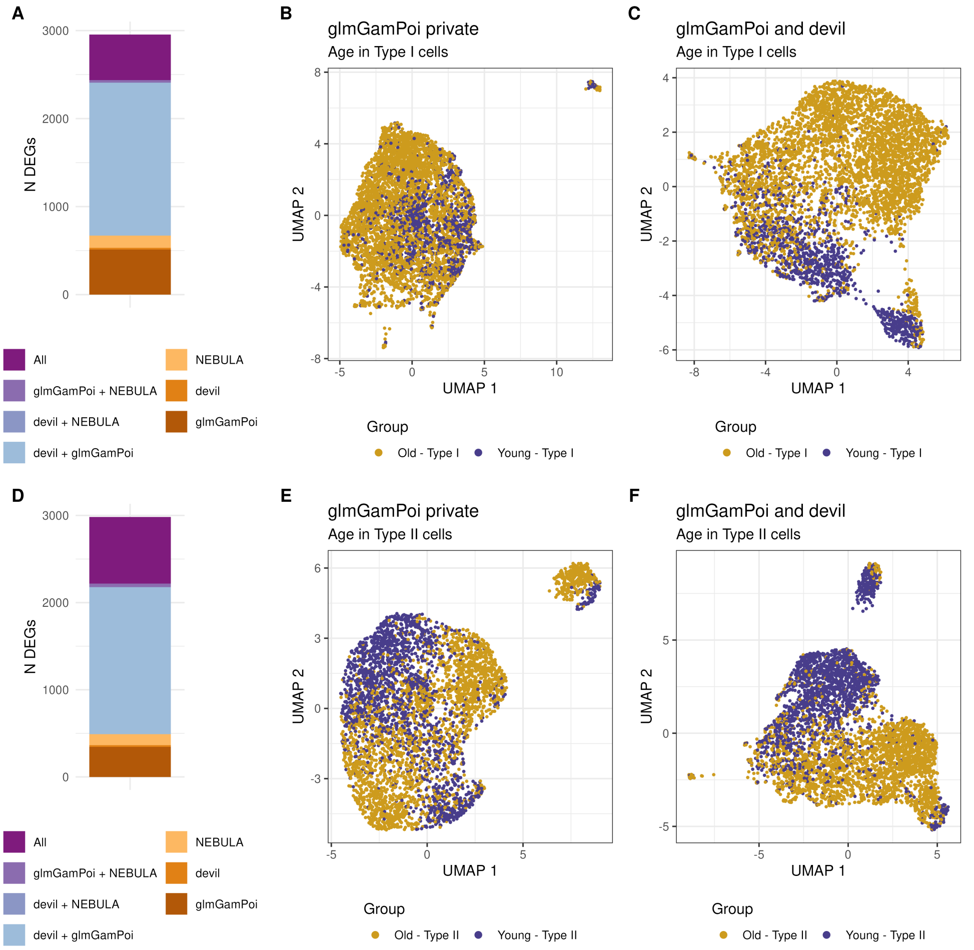


**Supplementary Figure 20 Impact of glmGamPoi-private and shared DE genes on ageing-related UMAP structure. A,D.** Stacked bar plots showing the number of differentially expressed genes (DEGs) identified uniquely by glmGamPoi, uniquely by devil, shared between methods, or detected by all methods, for Type I (A) and Type II (D) myonuclei. **B,E.** UMAP embeddings constructed using top 100 DE genes identified exclusively by glmGamPoi for ageing contrasts in Type I (B) and Type II (E) myonuclei. **C,F.** UMAP embeddings constructed using using top 100 DE genes shared between glmGamPoi and devil for ageing contrasts in Type I (C) and Type II (F) myonuclei.

### 2 Supplementary Methods

#### 2.1 Model description

##### 2.1.1 Modelling framework

Negative Binomial (NB) regression is a Generalised Linear Model (GLM) tailored to count data with overdispersion, which makes it a natural choice for RNA-seq. Given a count matrix $\mathbf{Y} \in \mathbb{R}^{N \times G}$ (with $n = 1, \ldots, N$ indexing cells and $g = 1, \ldots, G$ indexing genes) and a design matrix $\mathbf{X} \in \mathbb{R}^{N \times F}$ (with $f = 1, \ldots, F$ indexing covariates), the GLM is written as

$$\mathbb{E}[\mathbf{Y} \mid \mathbf{X}] \;=\; \boldsymbol{\mu} \;=\; g^{-1}(\mathbf{X}\boldsymbol{\beta}), \qquad \mathbf{Y} \sim \mathrm{NB}(\boldsymbol{\mu}, \boldsymbol{\theta}), \tag{1}$$

where $\boldsymbol{\beta} \in \mathbb{R}^{F \times G}$ is the matrix of regression coefficients, $\boldsymbol{\theta} \in \mathbb{R}^{G}$ the vector of overdispersion parameters, and $g$ is the link function mapping the linear predictor onto the support of the response mean.

Among the available parametrisations of the NB, we adopt

$$L(y \mid \mu, \theta) \;=\; \frac{\Gamma(\theta + y)}{y!\,\Gamma(\theta)} \left(\frac{\theta}{\theta + \mu}\right)^{\theta} \left(\frac{\mu}{\theta + \mu}\right)^{y}, \tag{2}$$

which yields

$$\mathbb{E}[Y] \;=\; \mu, \qquad \mathrm{Var}(Y) \;=\; \mu + \frac{\mu^2}{\theta}, \tag{3}$$

cleanly separating Poisson variance from biological overdispersion.

##### 2.1.2 Model likelihood

DEVIL requires: (i) a count matrix $\mathbf{Y} \in \mathbb{R}^{N \times G}$, where $y_{g,n}$ is the count of gene $g$ in cell $n$; (ii) a design matrix $\mathbf{X} \in \mathbb{R}^{N \times F}$, where $x_{n,f}$ is the value of feature $f$ for cell $n$; and, (iii) for multi-patient data, a patient-assignment vector $\mathbf{p} \in \mathbb{R}^{N}$. Raw counts are modelled as

$$y_{g,n} \;\sim\; \mathrm{NB}\big(\mu_{g,n} = s_n \,\exp\{\mathbf{x}_n \boldsymbol{\beta}_g\},\; \theta_g\big)\,, \tag{4}$$

where $\mathbf{x}_n = (x_{n,1}, \ldots, x_{n,F}) \in \mathbb{R}^{1 \times F}$ is the covariate row vector for cell $n$ and $\boldsymbol{\beta}_g = (\beta_{1,g}, \ldots, \beta_{F,g})^{\top} \in \mathbb{R}^{F \times 1}$ the gene-specific coefficient vector. The dispersion $\theta_g$ is shared across cells for a given gene, while the mean depends on the linear predictor through an exponential link.

The factor $s_n$ is a cell-specific size factor accounting for differences in sequencing depth across cells. It is computed prior to fitting using standard normalisation schemes (e.g. CPM, geometric-mean scaling, deconvolution- or Pareto-based methods [10–12]). We assume $s_{n,g} = s_n$, i.e. constant across genes for a given cell.

For inference, we exploit the equivalence of the NB distribution with a Poisson–Gamma mixture:

$$y_g \;\sim\; \mathrm{Poisson}(\lambda_g), \qquad \lambda_g \;\sim\; \mathrm{Gamma}\big(\theta_g,\, \theta_g \exp\{-\mathbf{X}\boldsymbol{\beta}_g\}\big)\,. \tag{5}$$

This representation is the basis of the variational inference scheme described below. DEVIL fits the GLM gene by gene in two steps: first $\boldsymbol{\beta}_g$ is estimated by Variational Inference, then $\theta_g$ is estimated by Maximum Likelihood (MLE) or Method of Moments (MoM).

#### Parameter inference

##### Variational posterior approximation

We consider a single gene and drop the subscript $g$ for clarity. The generative model is

$$\begin{aligned} \mathbf{y} \mid \boldsymbol{\lambda} &\sim \text{Poisson}(\boldsymbol{\lambda}), \\ \boldsymbol{\lambda} \mid \boldsymbol{\beta}, \theta &\sim \text{Gamma}(\theta,\, \theta \exp(-\mathbf{X}\boldsymbol{\beta}))\,, \\ \boldsymbol{\beta} &\sim \text{MVN}(\mathbf{0},\, \sigma_\beta^2\,\mathbf{I}), \\ \theta &\sim \delta(\theta - \theta_0), \end{aligned} \tag{6}$$

where the last line indicates that $\theta$ is held fixed at an initial estimate $\theta_0$. The joint distribution factorises as

$$P(\mathbf{y}, \boldsymbol{\lambda}, \boldsymbol{\beta}, \theta) \;=\; P(\boldsymbol{\beta})\, P(\boldsymbol{\lambda} \mid \boldsymbol{\beta}, \theta)\, P(\mathbf{y} \mid \boldsymbol{\lambda}), \tag{7}$$

with individual terms

$$\begin{aligned} P(\boldsymbol{\beta}) &= (2\pi)^{-F/2}\, |\Sigma_\beta|^{-1/2}\, \exp\Big\{-\tfrac{1}{2}\boldsymbol{\beta}^\top \Sigma_\beta^{-1} \boldsymbol{\beta}\Big\}\,, \\ P(\boldsymbol{\lambda} \mid \boldsymbol{\beta}, \theta) &= \prod_i \frac{\lambda_i^{\theta-1} \exp[-\theta \exp(-\mathbf{x}_i\boldsymbol{\beta})\lambda_i]\, [\theta \exp(-\mathbf{x}_i\boldsymbol{\beta})]^\theta}{\Gamma(\theta)}, \\ P(\mathbf{y} \mid \boldsymbol{\lambda}) &= \prod_i \frac{\lambda_i^{y_i} \exp(-\lambda_i)}{y_i!}. \end{aligned} \tag{8}$$

Equivalently, the log-terms read

$$\begin{aligned} \log P(\boldsymbol{\beta}) &= -\tfrac{F}{2}\log(2\pi\sigma_\beta^2) - \tfrac{1}{2}\boldsymbol{\beta}^\top \Sigma_\beta^{-1} \boldsymbol{\beta}, \\ \log P(\boldsymbol{\lambda} \mid \boldsymbol{\beta}, \theta) &= \sum_i [(\theta - 1)\log\lambda_i - \lambda_i\theta\exp(-\mathbf{x}_i\boldsymbol{\beta}) + \theta(\log\theta - \mathbf{x}_i\boldsymbol{\beta}) - \log\Gamma(\theta)]\,, \\ \log P(\mathbf{y} \mid \boldsymbol{\lambda}) &= \sum_i [y_i \log\lambda_i - \lambda_i - \log y_i!]\,. \end{aligned} \tag{9}$$

We approximate the posterior by a mean-field factorisation,

$$q(\boldsymbol{\beta}, \boldsymbol{\lambda} \mid \mathbf{y}) \;=\; q(\boldsymbol{\beta})\, q(\boldsymbol{\lambda}), \tag{10}$$

and update each factor via the standard coordinate-ascent rule

$$q_j^* \;\propto\; \exp\{\mathbb{E}_{-j}[\log p(z_j, \mathbf{z}_{-j}, \mathbf{x})]\}\,. \tag{11}$$

#### Variational update for $\boldsymbol{\lambda}$

Applying the rule to $\boldsymbol{\lambda}$, $\log q^*(\boldsymbol{\lambda}) \propto \mathbb{E}_{-\boldsymbol{\lambda}}[\log P(\mathbf{y}, \boldsymbol{\lambda}, \boldsymbol{\beta}, \theta)]$, and taking expectations under $q(\boldsymbol{\beta})$:

$$\begin{aligned}
&\mathbb{E}_{\theta,\boldsymbol{\beta},\mathbf{y}}[\log P(\boldsymbol{\beta}) + \log P(\boldsymbol{\lambda} \mid \boldsymbol{\beta}, \theta) + \log P(\mathbf{y} \mid \boldsymbol{\lambda})] \\
&\quad = \mathbb{E}\left[\sum_i (\theta - 1)\log \lambda_i - \lambda_i \theta \exp(-\mathbf{x}_i\boldsymbol{\beta}) + \theta(\log\theta - \mathbf{x}_i\boldsymbol{\beta}) - \log\Gamma(\theta) + y_i \log\lambda_i - \lambda_i - \log y_i!\right] \\
&\quad = \sum_i \left[(\theta + y_i - 1)\log\lambda_i - \lambda_i\big(1 + \theta\exp\{-\mathbf{x}_i\boldsymbol{\beta} + \tfrac{1}{2}\mathbf{x}_i\boldsymbol{\Sigma}_{\boldsymbol{\beta}}\mathbf{x}_i^\top\}\big)\right].
\end{aligned} \tag{12}$$

This is the kernel of a Gamma distribution with updated parameters

$$q^*(\boldsymbol{\lambda}) \;=\; \text{Gamma}\big(\theta + \mathbf{y},\, \mathbf{1} + \theta\exp\{-\mathbf{X}\boldsymbol{\beta} + \tfrac{1}{2}\,\text{diagonal}(\mathbf{X}\boldsymbol{\Sigma}_{\boldsymbol{\beta}}\mathbf{X}^\top)\}\big), \tag{13}$$

so at each iteration $\lambda_i$ is updated to the mean of that Gamma:

$$\lambda_i^* \;=\; \frac{\theta + y_i}{1 + \theta\exp\{-\mathbf{x}_i\boldsymbol{\beta} + \frac{1}{2}\mathbf{x}_i\boldsymbol{\Sigma}_{\boldsymbol{\beta}}\mathbf{x}_i^\top\}}. \tag{14}$$

#### Variational update for $\boldsymbol{\beta}$

We collect from $\mathbb{E}_q[\log p(\mathbf{y}, \boldsymbol{\lambda}, \boldsymbol{\beta}, \theta)]$ only the terms that depend on $\boldsymbol{\mu}_{\boldsymbol{\beta}} = \mathbb{E}_q[\boldsymbol{\beta}]$ or $\boldsymbol{\Sigma}_{\boldsymbol{\beta}}$:

$$\begin{aligned}
S \;\equiv\; & N\theta\log\theta - \theta\,\mathbf{1}^\top\mathbf{X}\boldsymbol{\mu}_{\boldsymbol{\beta}} - N\log\Gamma(\theta) + (\theta - 1)\,\mathbf{1}^\top\mathbb{E}_q[\log\boldsymbol{\lambda}] \\
& - \theta\,\mathbb{E}_q[\boldsymbol{\lambda}]^\top\,\mathbb{E}_q[\exp(-\mathbf{X}\boldsymbol{\beta})] \\
& - \tfrac{F}{2}\log(2\pi) - \tfrac{F}{2}\log(\sigma_\beta^2) - \tfrac{1}{2}\,\text{Tr}\Big(\sigma_\beta^{-2}\mathbf{I}\,\boldsymbol{\mu}_{\boldsymbol{\beta}}\boldsymbol{\mu}_{\boldsymbol{\beta}}^\top\Big).
\end{aligned} \tag{15}$$

The first two lines come from $\log P(\boldsymbol{\lambda} \mid \boldsymbol{\beta}, \theta)$, the last from $\log P(\boldsymbol{\beta})$. Using the moment-generating function of a Multivariate Normal, the middle term rewrites as

$$-\theta\,\mathbb{E}_q[\boldsymbol{\lambda}]^\top\,\exp\{-\mathbf{X}\boldsymbol{\mu}_{\boldsymbol{\beta}} + \tfrac{1}{2}\,\text{diagonal}(\mathbf{X}\boldsymbol{\Sigma}_{\boldsymbol{\beta}}\mathbf{X}^\top)\}. \tag{16}$$

To obtain the updates for $\boldsymbol{\mu}_{\boldsymbol{\beta}}$ and $\boldsymbol{\Sigma}_{\boldsymbol{\beta}}$, we use the result of Wand et al. [13], which states that

$$\begin{cases}
\boldsymbol{\Sigma}_{\boldsymbol{\beta}} \;\leftarrow\; \big\{-2\,\text{vec}^{-1}\big((D_{\text{vec}(\Sigma)}S)^\top\big)\big\}^{-1}, \\
\boldsymbol{\mu}_{\boldsymbol{\beta}} \;\leftarrow\; \boldsymbol{\mu}_{\boldsymbol{\beta}} + \boldsymbol{\Sigma}_{\boldsymbol{\beta}}\,D_{\boldsymbol{\mu}}S,
\end{cases} \tag{17}$$

where $\text{vec}(\mathbf{A})$ for a $d \times d$ matrix $\mathbf{A}$ denotes the $d^2 \times 1$ vector obtained by stacking the columns of $\mathbf{A}$.

The derivative with respect to $\boldsymbol{\mu}_{\boldsymbol{\beta}}$ is

$$\begin{aligned}
D_{\boldsymbol{\mu}}S &= -\theta\,\mathbf{X}^\top\mathbf{1} + \theta\,\mathbf{X}^\top\big[\mathbb{E}_q[\boldsymbol{\lambda}] \odot \exp\{-\mathbf{X}\boldsymbol{\mu}_{\boldsymbol{\beta}} + \tfrac{1}{2}\,\text{diagonal}(\mathbf{X}\boldsymbol{\Sigma}_{\boldsymbol{\beta}}\mathbf{X}^\top)\}\big] \\
&= \theta\,\mathbf{X}^\top\big[\mathbb{E}_q[\boldsymbol{\lambda}] \odot \exp\{-\mathbf{X}\boldsymbol{\mu}_{\boldsymbol{\beta}} + \tfrac{1}{2}\,\text{diagonal}(\mathbf{X}\boldsymbol{\Sigma}_{\boldsymbol{\beta}}\mathbf{X}^\top)\} - \mathbf{1}\big],
\end{aligned} \tag{18}$$

where $\odot$ denotes the Hadamard (elementwise) product. The derivative with respect to $\mathrm{vec}(\boldsymbol{\Sigma}_{\boldsymbol{\beta}})$ is

$$\begin{aligned} D_{\mathrm{vec}(\Sigma)} S &= D_{\mathrm{vec}(\Sigma)} \left[ -\theta\, \mathbb{E}_q[\boldsymbol{\lambda}]^\top \exp\{ -\mathbf{X}\boldsymbol{\mu}_{\boldsymbol{\beta}} + \tfrac{1}{2}\, \mathrm{diagonal}(\mathbf{X}\boldsymbol{\Sigma}_{\boldsymbol{\beta}}\mathbf{X}^\top) \} \right] \\ &= -\theta\, \mathbb{E}_q[\boldsymbol{\lambda}]^\top \,\mathrm{diag}\left[ \exp\{ -\mathbf{X}\boldsymbol{\mu}_{\boldsymbol{\beta}} + \tfrac{1}{2}\, \mathcal{Q}(\mathbf{X})\, \mathrm{vec}(\boldsymbol{\Sigma}_{\boldsymbol{\beta}}) \} \right] \tfrac{1}{2}\, \mathcal{Q}(\mathbf{X}) \\ &= -\tfrac{1}{2}\theta \left( \mathbb{E}_q[\boldsymbol{\lambda}] \odot \exp\{ -\mathbf{X}\boldsymbol{\mu}_{\boldsymbol{\beta}} + \tfrac{1}{2}\, \mathcal{Q}(\mathbf{X})\, \mathrm{vec}(\boldsymbol{\Sigma}_{\boldsymbol{\beta}}) \} \right)^\top \mathcal{Q}(\mathbf{X}) \\ &= -\tfrac{1}{2}\theta\, \mathrm{vec}\left( \mathbf{X}\, \mathrm{diag}\left[ \mathbb{E}_q[\boldsymbol{\lambda}] \odot \exp\{ -\mathbf{X}\boldsymbol{\mu}_{\boldsymbol{\beta}} + \tfrac{1}{2}\, \mathcal{Q}(\mathbf{X})\, \mathrm{vec}(\boldsymbol{\Sigma}_{\boldsymbol{\beta}}) \} \right] \mathbf{X} \right)^\top , \end{aligned} \tag{19}$$

where, for a $d \times 1$ vector $\mathbf{a}$, $\mathrm{diag}(\mathbf{a})$ denotes the $d \times d$ diagonal matrix with $\mathbf{a}$ on the diagonal. The derivation uses twice the identities of Wand et al. [13]: given an $n \times d$ matrix $\mathbf{A}$, a $d \times d$ matrix $\mathbf{B}$, and a $d \times 1$ vector $\mathbf{b}$, defining $\mathcal{Q}(\mathbf{A}) \equiv (\mathbf{A} \otimes \mathbf{1}^\top) \odot (\mathbf{1}^\top \otimes \mathbf{A})$ with $\otimes$ the Kronecker product, then

$$\mathrm{diag}(\mathbf{A}\mathbf{B}\mathbf{A}^\top) = \mathcal{Q}(\mathbf{A})\, \mathrm{vec}(\mathbf{B}), \tag{20}$$

$$\mathrm{vec}(\mathbf{A}^\top \mathrm{diag}(\mathbf{b})\mathbf{A}) = \mathcal{Q}(\mathbf{A})^\top \mathbf{b}. \tag{21}$$

Putting everything together, the variational update scheme is

$$\begin{cases} \boldsymbol{\Sigma}_{\boldsymbol{\beta}} \;\leftarrow\; \left\{ \theta\, \mathbf{X}^\top \mathrm{diag}\left[ \mathbb{E}_q[\boldsymbol{\lambda}] \odot \exp\{ -\mathbf{X}\boldsymbol{\mu}_{\boldsymbol{\beta}} + \tfrac{1}{2}\, \mathrm{diagonal}(\mathbf{X}\boldsymbol{\Sigma}_{\boldsymbol{\beta}}\mathbf{X}^\top) \} \right] \mathbf{X} \right\}^{-1}, \\ \boldsymbol{\mu}_{\boldsymbol{\beta}} \;\leftarrow\; \boldsymbol{\mu}_{\boldsymbol{\beta}} + \boldsymbol{\Sigma}_{\boldsymbol{\beta}} \left\{ \theta\, \mathbf{X}^\top \left[ \mathbb{E}_q[\boldsymbol{\lambda}] \odot \exp\{ -\mathbf{X}\boldsymbol{\mu}_{\boldsymbol{\beta}} + \tfrac{1}{2}\, \mathrm{diagonal}(\mathbf{X}\boldsymbol{\Sigma}_{\boldsymbol{\beta}}\mathbf{X}^\top) \} - \mathbf{1} \right] \right\}. \end{cases} \tag{22}$$

The full algorithm, including initialisation ($\beta_0 = \log(1 + \bar{y}_g)$ for the intercept, zero otherwise) and the convergence criterion, is given in Supplementary Algorithms 1 and 2.

#### 2.2 Overdispersion inference

Once $\hat{\boldsymbol{\beta}}_g$ is obtained, $\theta_g$ is estimated conditional on the fitted means $\mu_{g,i} = s_i \exp\{\mathbf{x}_i \hat{\boldsymbol{\beta}}_g\}$. DEVIL supports two estimators.

##### 2.2.1 Maximum-likelihood estimator (MLE)

The NB log-likelihood for gene $g$ is

$$\begin{aligned} \ell(\theta_g) = \sum_{i=1}^{N} \Big[ & \log\Gamma(y_{g,i} + \theta_g) - \log\Gamma(\theta_g) - \log(y_{g,i}!) \\ & + \theta_g \log\left( \tfrac{\theta_g}{\theta_g + \mu_{g,i}} \right) + y_{g,i} \log\left( \tfrac{\mu_{g,i}}{\theta_g + \mu_{g,i}} \right) \Big], \end{aligned} \tag{23}$$

and the MLE is

$$\hat{\theta}_g^{\mathrm{MLE}} = \arg\max_{\theta_g > 0} \ell(\theta_g), \tag{24}$$

obtained by iterative numerical optimisation. The score function involves the digamma function $\psi(\cdot)$ and the Hessian the trigamma function $\psi_1(\cdot)$, which makes efficient GPU execution impractical.

##### 2.2.2 Method-of-moments estimator (MoM)

Defining the residuals $r_{g,i} = y_{g,i} - \mu_{g,i}$ and approximating

$$\mathrm{Var}(Y_{g,i}) \;\approx\; \frac{1}{N-1}\sum_{i=1}^{N} r_{g,i}^2 \;=\; \mu_{g,i} + \frac{\mu_{g,i}^2}{\theta_g}, \tag{25}$$

yields the MoM estimator

$$\hat{\theta}_g^{\mathrm{MoM}} \;=\; \left[\frac{\sum_i (\mu_{g,i} - y_{g,i})^2 - \sum_i \mu_{g,i}}{\sum_i \mu_{g,i}^2}\right]_+^{-1}, \tag{26}$$

where $[\,\cdot\,]_+$ denotes truncation to positive values. Unlike the MLE, this estimator requires only elementwise operations and reductions, making it fully GPU-compatible. It is less efficient than the MLE in small samples but well suited to large-scale analyses.

#### 2.3 Covariance structure estimation

The variational posterior covariance $\boldsymbol{\Sigma}_{\boldsymbol{\beta}}$ obtained during inference systematically underestimates uncertainty: it inherits the bias of the mean-field assumption and, critically, does not incorporate $\theta_g$ into the variance structure. Using it directly for testing produces underestimated standard errors and inflated test statistics, even in the single-patient case. DEVIL therefore re-estimates $\boldsymbol{\Sigma}_{\boldsymbol{\beta}}$ from the fitted parameters after inference, with two strategies depending on the experimental design.

In what follows, $l = \log p(\mathbf{y}, \mathbf{X}, \boldsymbol{\beta}, \theta)$ denotes the NB log-likelihood from equation (2), and we work with a generic IID sample to derive the general result before specialising to our setting. The exposition follows the classical sandwich-estimator framework, which we summarise here for completeness.

##### 2.3.1 Correctly specified IID model

Consider IID random variables $\mathbf{x} = (x_1, \ldots, x_N)$ drawn from a density $L_{\boldsymbol{\beta}}$ indexed by the parameter vector $\boldsymbol{\beta}$. The joint likelihood and log-likelihood are

$$L_{N,\boldsymbol{\beta}}(\mathbf{x}) = \prod_{i=1}^{N} L_{\boldsymbol{\beta}}(x_i), \qquad l_{N,\boldsymbol{\beta}}(\mathbf{x}) = \sum_{i=1}^{N} \log L_{\boldsymbol{\beta}}(x_i). \tag{27}$$

If the model is correctly specified, differentiating $\int L_{\boldsymbol{\beta}}(x)\,dx = 1$ yields the two classical identities

$$\mathbb{E}_{\boldsymbol{\beta}}[l'(\boldsymbol{\beta})] \;=\; 0, \tag{28a}$$

$$\mathrm{Var}_{\boldsymbol{\beta}}[l'(\boldsymbol{\beta})] \;=\; -\mathbb{E}_{\boldsymbol{\beta}}[l''(\boldsymbol{\beta})], \tag{28b}$$

both equal to the Fisher information. We denote by $I_N(\boldsymbol{\beta})$ the Fisher information for a sample of size $N$, and by $I_1(\boldsymbol{\beta})$ its per-observation version. Applying the central limit theorem to the empirical score gives

$$\sqrt{N}\,\frac{1}{N}\,l'_N(\boldsymbol{\beta}) \;\xrightarrow{D}\; \mathcal{N}(\mathbf{0},\, I_1(\boldsymbol{\beta})), \tag{29}$$

and, similarly,

$$-\frac{1}{N}\,l''_N(\boldsymbol{\beta}) \;\xrightarrow{P}\; I_1(\boldsymbol{\beta}). \tag{30}$$

For the MLE $\hat{\boldsymbol{\beta}}$, defined by $l'_N(\hat{\boldsymbol{\beta}}) = 0$, a Taylor expansion around the true $\boldsymbol{\beta}$ gives

$$0 \;=\; l'_N(\hat{\boldsymbol{\beta}}) \;\approx\; l'_N(\boldsymbol{\beta}) + l''_N(\boldsymbol{\beta})(\hat{\boldsymbol{\beta}} - \boldsymbol{\beta}), \tag{31}$$

and combining with equations (29) and (30) yields the well-known asymptotic distribution

$$\sqrt{N}(\hat{\boldsymbol{\beta}} - \boldsymbol{\beta}) \;\xrightarrow{D}\; \mathcal{N}\big(\mathbf{0},\, I_1(\boldsymbol{\beta})^{-1}\big)\,. \tag{32}$$

##### 2.3.2 Misspecified model

When the model is not correctly specified, i.e. the data are generated by a density $h$ that is not in the family $\{L_{\boldsymbol{\beta}}\}$, equations (28a) and (28b) no longer hold. Defining

$$\lambda_h(\boldsymbol{\beta}) \;=\; \mathbb{E}_h[l(\boldsymbol{\beta})] \tag{33}$$

and assuming $\lambda_h$ has a maximum at $\boldsymbol{\beta}^*$, so that $\mathbb{E}_h[l'(\boldsymbol{\beta}^*)] = 0$, we introduce

$$M(\boldsymbol{\beta}) \;=\; \mathrm{Var}_h[l'(\boldsymbol{\beta})] \qquad \text{(meat)}, \tag{34a}$$

$$B(\boldsymbol{\beta}) \;=\; \big(-\mathbb{E}_h[l''(\boldsymbol{\beta})]\big)^{-1} \qquad \text{(bread)}. \tag{34b}$$

These coincide with $I_1(\boldsymbol{\beta})$ and $I_1(\boldsymbol{\beta})^{-1}$ respectively under correct specification. Note that we have absorbed the inverse into the definition of $B$ so the sandwich appears in its simplest form below. Repeating the central-limit argument under $h$,

$$\sqrt{N}\,\frac{1}{N}\,l'_N(\boldsymbol{\beta}^*) \;\xrightarrow{D}\; \mathcal{N}(\mathbf{0},\, M(\boldsymbol{\beta}^*)), \qquad -\frac{1}{N}\,l''_N(\boldsymbol{\beta}^*) \;\xrightarrow{P}\; B(\boldsymbol{\beta}^*)^{-1}, \tag{35}$$

leads to the asymptotic distribution

$$\sqrt{N}(\hat{\boldsymbol{\beta}} - \boldsymbol{\beta}^*) \;\xrightarrow{D}\; \mathcal{N}(\mathbf{0},\, B(\boldsymbol{\beta}^*)\,M(\boldsymbol{\beta}^*)\,B(\boldsymbol{\beta}^*))\,. \tag{36}$$

This is the so-called sandwich $BMB$, with $B$ playing the role of the two slices and $M$ the filling [14–17].

In practice, $B$ and $M$ are estimated from data as

$$\hat{B} \;=\; \left(\frac{1}{N}\sum_{i=1}^{N} -l''(y_i \mid x_i, \boldsymbol{\beta}, \theta)\right)^{-1}, \qquad \hat{M} \;=\; \frac{1}{N}\sum_{i=1}^{N} l'(y_i \mid x_i, \boldsymbol{\beta}, \theta)\, l'(y_i \mid x_i, \boldsymbol{\beta}, \theta)^{\top}. \tag{37}$$

##### 2.3.3 Single-patient case

When all cells come from a single patient, $\boldsymbol{\Sigma}_{\boldsymbol{\beta}}$ is the inverse of the negative empirical Hessian:

$$\boldsymbol{\Sigma}_{\boldsymbol{\beta}} \;=\; -H_l^{-1}, \qquad H_l \;=\; \frac{1}{N}\sum_{i=1}^{N} l''(y_i, \mathbf{x}_i, \boldsymbol{\beta}, \theta), \tag{38}$$

where $l''$ is the second derivative of the NB log-likelihood with respect to $\boldsymbol{\beta}$.

##### Multi-patient case (clustered sandwich estimator)

When cells are nested within patients, the IID assumption fails: cells from the same patient are correlated, and the inverse-Hessian underestimates variance by ignoring intra-patient correlation. The bread $B$ is still given by equation (34b), but the meat $M$ must be replaced by a clustered version that sums scores within each patient before forming the outer product:

$$M_{CL} \;=\; \frac{1}{N}\sum_{p=1}^{P}\left[\sum_{i\in\mathcal{I}_p} l'(y_{i,g} \mid x_{i,g}, \boldsymbol{\beta}, \theta)\right]\left[\sum_{i\in\mathcal{I}_p} l'(y_{i,g} \mid x_{i,g}, \boldsymbol{\beta}, \theta)\right]^{\top}, \tag{39}$$

where $P$ is the number of patients and $\mathcal{I}_p$ the index set of cells from patient $p$. This estimator allows arbitrary correlation within each patient while still assuming independence between patients. The clustered covariance is then

$$\boldsymbol{\Sigma}_{\boldsymbol{\beta}} \;=\; B\, M_{CL}\, B. \tag{40}$$

##### 2.3.4 Effective sample size interpretation.

To see why $M_{CL}$ inflates variance, consider the scalar case and let $r_{i,g} = l'(y_{i,g} \mid x_{i,g}, \boldsymbol{\beta}, \theta)$. Then

$$M \;\propto\; \sum_i r_{i,g}^2, \qquad M_{CL} \;\propto\; \left(\sum_i r_{i,g}\right)^2 \;=\; \sum_i r_{i,g}^2 + 2\sum_{i<j} r_{i,g}\, r_{j,g}, \tag{41}$$

so $M_{CL} = M + \alpha$, with $\alpha > 0$ whenever within-patient scores are positively correlated. This inflation propagates through the sandwich formula, widening confidence intervals and producing more conservative p-values. Equivalently, the adjustment can be read as replacing the nominal sample size $N$ with an effective sample size $n_{\text{eff}} < N$, reflecting the reduced amount of independent information in nested data. Although the nominal number of observations is $N$, the presence of intra-patient correlation means the information content is lower, equivalent to having fewer, independent samples. As a result, standard errors increase and statistical tests become more conservative. This correction is essential to control false positives in the hierarchical structures typical of single-cell studies.

#### 2.4 Differential expression testing

With $\hat{\boldsymbol{\beta}}_g$ and $\hat{\boldsymbol{\Sigma}}_{\boldsymbol{\beta},g}$ estimated for each gene, the inference of $\boldsymbol{\beta}_g$ together with its covariance matrix allows hypothesis testing on linear combinations of coefficients. Following standard practice, DEVIL tests null hypotheses of the form $H_0 : \mathbf{c}^\top \boldsymbol{\beta}_g = 0$ via the Wald statistic

$$W_g = (\mathbf{c}^\top \hat{\boldsymbol{\beta}}_g)^\top \left[\mathbf{c}^\top \hat{\boldsymbol{\Sigma}}_{\boldsymbol{\beta},g}\, \mathbf{c}\right]^{-1} (\mathbf{c}^\top \hat{\boldsymbol{\beta}}_g) \xrightarrow{D} \chi^2, \tag{42}$$

exploiting the fact that $\hat{\boldsymbol{\beta}}_g$ is asymptotically Multivariate Normal with covariance $\hat{\boldsymbol{\Sigma}}_{\boldsymbol{\beta},g}$. When multiple genes are tested simultaneously, the resulting p-values are corrected across genes with the Benjamini–Hochberg procedure [18] to control the false discovery rate. This hybrid design, variational inference for scalable parameter estimation, classical robust standard errors for testing, combines computational efficiency with statistical rigour.

#### 2.5 GPU implementation details

##### 2.5.1 $\boldsymbol{\beta}$ coefficient initialisation and inference

The $\beta$ fitting procedure is arithmetically intensive and requires minimal branching, making it well-suited for GPU acceleration. The variational update equations are reformulated as three-dimensional tensor contractions using the cuTENSOR library[19]. This reformulation serves two purposes: it replaces multiple Level-1 and Level-2 BLAS calls with a single high-throughput kernel, and it allows cuTENSOR to optimise memory access patterns via coalesced reads, shared memory reuse, and register allocation. Mixed-precision arithmetic is supported natively; we select 32-bit floating-point for input and output tensors, promoting to 64-bit for intermediate accumulations where required to avoid numerical instability, consistent with the CPU reference implementation.

At each variational iteration, computing the covariance update $\Sigma_\beta$ requires inverting a batch of small matrices of size $F \times F$(where $F$ is the number of model covariates; typically $F < 10$ in practice). To minimise the per-inversion kernel-launch overhead, we use the batched LU factorisation with partial pivoting and batched matrix inversion routines from cuBLAS[20]. These routines include specialised kernels for small matrix dimensions, selected automatically based on data type and matrix size. Performance was confirmed via GPU profiling, which validated the use of size-specialised kernels for the matrix dimensions encountered in typical devil analyses.

Coefficients are initialised by setting the intercept term to $\log(1 + \bar{y}_g)$ and all remaining coefficients to zero. This initialisation requires only row-wise averaging of the input count matrix, a memory-bandwidth-bound operation that is highly efficient on GPU hardware.

The full GPU pseudocode for $\beta$ inference is provided in Supplementary Algorithm 4.

##### 2.5.2 Overdispersion inference

Following convergence of $\beta$, the MoM overdispersion estimator is computed entirely on-device using cuBLAS for Level-2 and Level-3 BLAS operations (matrix-vector products, reductions) and custom CUDA kernels for element-wise computations (residual calculation, truncation to positive values). No CPU round-trip is required between $\beta$ and $\theta$ inference steps. The full GPU pseudocode for $\theta$ inference is provided in Supplementary Algorithm 3.

##### 2.5.3 Multi-GPU scaling.

Multi-GPU execution is implemented by spawning one CPU thread per GPU device. Each thread processes an independent batch of genes, with batches distributed across devices at runtime. Batch size controls the trade-off between parallelisation overhead (smaller batches reduce load imbalance from variable per-gene convergence) and launch overhead (larger batches amortise kernel-launch costs). A batch size of 100 genes was found empirically to provide the best balance across these factors while keeping per-device memory consumption below 10 GB, compatible with standard HPC GPU nodes.

##### 2.5.4 Software and hardware environment.

All benchmarks were performed on the ORFEO HPC cluster. GPU experiments used nodes with either 8× NVIDIA A100 (40 GB HBM2) or 8× NVIDIA H100 (80 GB HBM3) accelerators. CPU benchmarks were run on the H100 nodes (same hardware) to ensure a fair comparison of CPU-only performance. The software stack comprised: CUDA Toolkit 12.6, cuBLAS 12.6, cuTENSOR 2.2.0.0, R 4.3.3, and OpenBLAS 0.3.29. All components — including devil, R, and OpenBLAS — were compiled from source directly on the target architecture to maximise instruction-level optimisation. A full description of node specifications and memory configurations is provided in Supplementary Table 10.

#### 3 Pseudocode and algorithm

**Algorithm 1 -** $\boldsymbol{\beta}_g$ Initialization

**Require:** $\mathbf{y}_g$, $C \in \mathbb{N}(Cells)$
  $\boldsymbol{\eta} \leftarrow \frac{1}{C} \sum_i \mathbf{y}_g[i]$
  $\boldsymbol{\beta}_g \leftarrow \mathbf{0}$
  $\boldsymbol{\beta}_g[0] \leftarrow \log(\mathbf{1} + \boldsymbol{\eta})$

**Algorithm 2 -** $\boldsymbol{\beta}_g$ inference

**Require:** $\mathbf{y}_g, \boldsymbol{\beta}_g, \mathbf{X},$ initialized$, \theta \geq 0, \epsilon \geq 0$
  $\boldsymbol{\Sigma}_{\boldsymbol{\beta}} \leftarrow \mathbf{I}$
  $\delta \leftarrow \infty$
  **while** norm($\delta$) $\geq \epsilon$ **do**
    $\mathbf{w} \leftarrow \exp\{-\mathbf{X}^T\boldsymbol{\beta}_g + \frac{1}{2}\text{diagonal}(\mathbf{X}\boldsymbol{\Sigma}_\beta\mathbf{X}^T)]\}^{-1}$
    $\lambda \leftarrow \frac{\mathbf{y}_g+\theta}{\mathbf{w}+\theta}$
    $\boldsymbol{\Sigma}_{\boldsymbol{\beta}} \leftarrow \{\mathbf{X}^T\text{diag}[\lambda \odot \mathbf{w}]\mathbf{X}\}^{-1}$
    $\delta \leftarrow \boldsymbol{\Sigma}_{\boldsymbol{\beta}}\mathbf{X}^T(\lambda \odot \mathbf{w} - \mathbf{1})$
    $\boldsymbol{\beta}_g \leftarrow \boldsymbol{\beta}_g + \delta$
  **end while**

**Algorithm 3 -** $\boldsymbol{\theta}$ estimation, Method of Moments (MoM)

**Require:** $\mathbf{y}_g, \mathbf{X}, \boldsymbol{\beta}_g, \mathbf{o}$
  $\boldsymbol{\eta} \leftarrow \boldsymbol{\beta}_g\mathbf{X}^T + \mathbf{o}$
  $\boldsymbol{\mu} \leftarrow \exp(\boldsymbol{\eta})$
  $\mathbf{N} \leftarrow (\mathbf{y} - \boldsymbol{\mu})^2 - \boldsymbol{\mu}$
  $\mathbf{D} \leftarrow \boldsymbol{\mu}^2$
  $\rho \leftarrow \frac{C}{C-F}$    ▷ $C$: cell count, $F$: feature count
  $\boldsymbol{\theta} \leftarrow \rho \cdot \frac{\mathbf{N}}{\mathbf{D}}$
  $\boldsymbol{\theta} \leftarrow \max(\boldsymbol{\theta}, 0)$

**Algorithm 4** - Batched $\boldsymbol{\beta}$ inference with GPU Acceleration, it runs on $N$ GPUs in parallel

**Input:** Input matrix: $Y \in \mathbb{R}^{G\times N}$, Feature matrix: $X \in \mathbb{R}^{F\times C}$, offset vector: offset $\in \mathbb{R}^{C}$, $k$, $\mu_\beta$, convergence tolerance: $\varepsilon$, max iterations max_iter
**Output:** Updated $\mu_\beta$ parameters for all gene batches
**Copy** $X[0{:}F,\ 0{:}C]$ to GPU ▷ F = features, C = cells, G = genes
**Copy** offset$[0{:}C]$ to GPU
**for each batch** of $G$ genes **do**
  **Copy** $X[batch,\ 0{:}C]$ to GPU ▷ $batch$ = batch indexes
  **Copy** $k[batch]$ and $\mu_\beta[batch]$ to GPU
  Initialize norm $\leftarrow \varepsilon + 1$, iter $\leftarrow 0$
  **while** iter $<$ max_iter **and** norm $> \varepsilon$ **do**
    iter $\leftarrow$ iter + 1
    cg_tmp $\leftarrow$ gpu_einsum$(ik, jk \rightarrow ji,\ X,\ \mu_\beta)$
    $w_q \leftarrow$ GPUKernel $\left[\exp\left(\text{cg\_tmp}_{i,j} - \text{offset}_i\right)\right]$ $\forall i \in [1, C],\ \forall j \in [1, G]$
    $\mu_g[i] \leftarrow$ GPUKernel $\left[\frac{k[i]+Y[i,j]}{1+k[i]\cdot w_q[i,j]}\right]$ $\forall i, j$
    $\mu_g[i] \leftarrow$ GPUKernel $[\mu_g[i] \cdot w_g[i]]$ $\forall i$
    $A \leftarrow$ gpu_einsum$(cf, gc \rightarrow cfg,\ X,\ \mu_g)$
    $B \leftarrow$ gpu_einsum$(cfg, ck \rightarrow gkf,\ A,\ X)$
    $B_k \leftarrow$ gpu_einsum$(gfc, g \rightarrow gfc,\ B,\ k)$
    $\Sigma \leftarrow$ Inverse$(B_k)$ ▷ LU decomposition with pivoting
    $\mu_g[i] \leftarrow$ GPUKernel $[\mu_g[i] - 1]$ $\forall i$
    $C \leftarrow$ gpu_einsum$(cf, gc \rightarrow gf,\ X,\ \mu_g)$
    $D \leftarrow$ gpu_einsum$(g, gf \rightarrow gf,\ k,\ C)$
    $\delta \leftarrow$ gpu_einsum$(gfk, gk \rightarrow gf,\ \Sigma,\ D)$
    $\mu_\beta[i] \leftarrow$ GPUKernel $[\mu_\beta[i] + \delta[i]]$ $\forall i$
    norm $\leftarrow$ MaxAbs$(\delta)$
  **end while**
  **Copy** updated $\mu_\beta[batch]$ to HOST
**end for**

[19] NVIDIA Corporation: cuTENSOR Documentation. https://docs.nvidia.com/cuda/cutensor/

[20] NVIDIA Corporation: cuBLAS Documentation. https://docs.nvidia.com/cuda/cublas/
